# Sequence adaptations satisfy the constraints of mitochondrial membrane protein evolution

**DOI:** 10.64898/2026.08.04.742770

**Authors:** Tarun Yadav, Jonathan H. Borowsky, Alvaro Jesus Narbona-Perez, Bhavya Soni, Corey N. Cunningham, James Carrington, Michael Grabe, Jared Rutter

**Affiliations:** Department of Biochemistry, University of Utah, Salt Lake City, United States; Cardiovascular Research Institute, Department of Pharmaceutical Chemistry, University of California, San Francisco, United States; Howard Hughes Medical Institute, University of Utah, Salt Lake City, United States

## Abstract

Inner mitochondrial membrane proteins must be sufficiently hydrophilic to withstand aqueous exposure during translation and transit to the mitochondria. Meanwhile, their transmembrane segments must be sufficiently hydrophobic to stably embed in the lipid membrane. We hypothesized that sequence-level adaptations evolved to balance these constraints. Here, we integrate structure-informed evolutionary analyses of mitochondrial proteins with atomistic simulations and cell-based experiments to identify aliphatic-to-threonine substitutions (ATS) as a potential solution to these constraints. With high statistical confidence, this transmembrane segment-specific adaptation is recurrently and convergently observed throughout mitochondrial evolution. Conformational analyses show that threonine interacts with both water and the transmembrane helix backbone, thereby lowering hydrophobicity without destabilizing secondary structure. In the extremely hydrophobic ATP6 protein, reverting threonines to aliphatic residues disrupts mitochondrial targeting, while introducing threonines into a poorly targeted variant improves its mitochondrial localization. These findings have implications for mitochondrial genome evolution, the rational design of membrane proteins, and potentially mitochondrial gene therapy.

## INTRODUCTION

Mitochondria execute numerous cellular processes, many of which rely on the proteins of the inner mitochondrial membrane (IMM) that play outsized roles in energy metabolism, macromolecule biosynthesis, and metabolite transport (Spinelli and Haigis, 2018). The mitochondrial proteome is encoded by two genomes. The mitochondrial DNA (mtDNA) resides in the mitochondrial matrix and encodes a small number of proteins, typically hydrophobic respiratory proteins that are embedded in the inner mitochondrial membrane. The vast majority of mitochondrial proteins are encoded by the nuclear genome, translated on cytosolic ribosomes, and post-translationally translocated into the mitochondria (Gray, 2015). Hence, mitochondrial function requires proteins whose biogenesis must be spatially and temporally coordinated across genomes, translation machineries, and insertion pathways (Soto et al., 2022).

The biogenesis of mtDNA-encoded IMM proteins, which are typically hydrophobic and contain multi-pass transmembrane domains, requires accurate folding and assembly into multi-subunit complexes. Specialized machinery exists to support this process including chaperones and insertases (Poerschke et al., 2024; Vercellino and Sazanov, 2022). Co-translational membrane insertion by the Oxa1L insertase may not shield TM helices (TMHs) from aqueous exposure to the same extent as bacterial and endoplasmic reticulum insertion machineries do (Figure 1A, 1B) (Itoh et al., 2021; Mercier et al., 2022; Schöndorf et al., 2026). Even more challenging aqueous exposure occurs for nuclear-encoded IMM proteins, which must traverse an aqueous environment prior to IMM incorporation (Nowicka et al., 2021).

**Figure 1:**
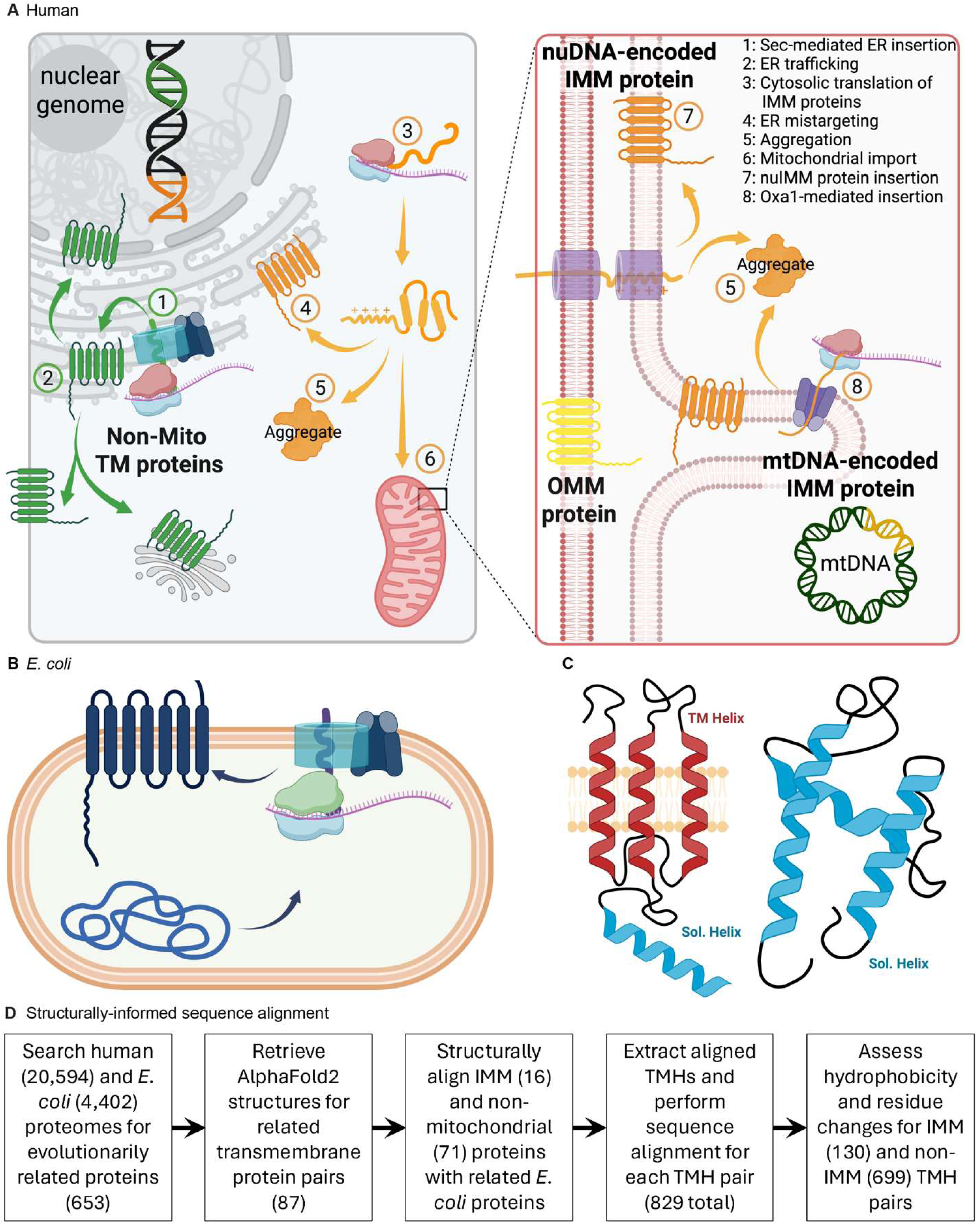
Mitochondrial transmembrane proteins face a solubility-stability tradeoff. **(A)** Non-mitochondrial proteins are co-translationally inserted via the SRP–Sec61 pathway with accessory Oxa1-like insertases (e.g. EMC/GET/TMCO) — bypassing solubility constraints (1, 2). Nuclear-encoded IMM proteins are post-translationally targeted (3), remaining vulnerable to mistargeting (4) and aggregation in the cytoplasm or during mitochondrial import (5). Once successfully imported (6), nuDNA-encoded IMM proteins are inserted into the IMM via TIM machinery and can be assisted by OXA1L (7). mtDNA-encoded IMM proteins, by contrast, are co-translationally inserted via OXA1L directly from the matrix (8), but face hydrophobicity-driven assembly and proteostasis constraints (5). **(B)** Bacterial (*E. coli*) proteins are co-translationally inserted into the cell membrane via the SRP– Sec pathway with an Oxa1-like insertase (YidC), bypassing aqueous exposure and proteostasis constraints. **(C)** Schematic illustrating transmembrane (red) and soluble (blue) helix categories analyzed from TM proteins (left) and soluble proteins (right). **(D)** Computational pipeline for structural alignment of evolutionarily related TMHs between *E. coli* and *H. sapiens*.

This demand for solubility in an aqueous environment conflicts with the need for integral IMM proteins to reside stably in the hydrophobic core of the IMM, which requires their TMHs to be enriched in aliphatic residues (Hessa et al., 2005). Thus, mitochondrial TM proteins face a biophysical tradeoff: they must be hydrophobic enough to insert and remain stable in the IMM, yet sufficiently compatible with aqueous environments to avoid aggregation during biogenesis. This tradeoff is particularly consequential in mitochondria, where the biogenesis machinery is comparatively streamlined (Hegde and Keenan, 2022; Tong et al., 2011), and chaperone systems are believed to operate near saturation (Banerjee et al., 2026; Preissler et al., 2015). Recent efforts to understand this paradox have examined the translocation machineries, chaperone mechanisms, and the roles of mitochondrial targeting sequences (MTSs) in subcellular localization (Juszkiewicz et al., 2025). In contrast, we hypothesized that adaptations at the level of individual protein sequences could be revealed by examining mitochondrial proteins across evolutionary time.

Here, we identify aliphatic-to-threonine substitutions (ATS) as a recurrent and TMH-specific feature of IMM proteins that addresses this biochemical challenge. Combining structure-informed comparative genomics spanning evolutionarily diverse proteomes, molecular dynamics simulations, and cell-based experiments, we show that mitochondrial TMHs are selectively enriched in threonine at the expense of aliphatic residues. These results show that ATS can lower mitochondrial TMH hydrophobicity while preserving IMM compatibility. In cells, native ATS is required for efficient localization of an IMM protein, and engineered ATS can improve localization of an otherwise mistargeted IMM protein. We further demonstrate that the ATS signature is also recovered in IMM proteins that underwent mtDNA-to-nuDNA gene transfer events, supporting ATS as an effective strategy for resolving the solubility-stability tradeoff over mitochondrial evolution. Together, our findings offer a sequence-level view of the biochemical constraints shaping mitochondrial proteome evolution and how ATS could resolve them.

## RESULTS

### Evolutionary comparisons show aliphatic-to-threonine substitutions in mitochondrial transmembrane proteins

To test whether IMM proteins exhibit adaptations to address the solubility-stability tradeoff, we compared human TM proteins with evolutionarily related proteobacterial proteins lacking similar biogenesis challenges. We used the *E. coli* proteome because of its comprehensive annotation and the availability of structural information (Figure 1A, B). For comparing highly divergent proteins, traditional primary sequence-based methods can be unreliable and produce large mapping gaps and misaligned regions (Carpentier and Chomilier, 2019; Rajapaksa et al., 2023). We therefore used structurally informed sequence alignment, which enabled robust one-to-one pairwise comparison of TMHs within each protein pair (Figure 1D, Figure 1—figure supplement 1A–C) — a prerequisite for the residue-level analyses that follow.

Using this dataset (supplementary table A), we assessed the compositional differences between paired *E. coli* and human TMHs (Figure 1C). Compared to related proteobacterial proteins, both nuDNA- and mtDNA-encoded human mitochondrial proteins showed reduced TMH hydrophobicity (Figure 2A, B). To determine how this reduced hydrophobicity is accomplished while maintaining membrane stability, we analyzed the relative amino acid composition of human vs. *E. coli* TMHs (Figure 2C), and we found that the decrease in hydrophobicity is driven primarily by a reduction in aliphatic residues (A, L, I, V, F, hereafter ALIVF) (Figure 2D, E). In parallel, we found a significant enrichment of threonine (T) residues in human mitochondrial TMHs regardless of whether they are encoded by the nuclear or mitochondrial genome (Figure 2C, F, G). By contrast, TMHs of human proteins not localized to the IMM exhibited a very minor decrease in hydrophobicity relative to paired *E. coli* TMHs and no increase in threonine content (Figure 2— figure supplement 1A, B). Instead, non-mitochondrial TMHs were enriched for serine rather than threonine residues, despite sharing the same genomic origin as nuclear-encoded mitochondrial TMHs (Figure 2—figure supplement 1C). We did not observe these changes when similar analyses were carried out for soluble helices from the same TM proteins (Figure 1C, Figure 2—figure supplement 1D–F), nor with fully soluble mitochondrial proteins (Figure 1C, Figure 2—figure supplement 2, supplementary table B). This suggests that these observations of reduced ALIVF content and increased threonine content in mitochondrial proteins are not due to pan-mitochondrial confounders — such as the protein import machinery or adaptation to the mitochondrial environment.

**Figure 2:**
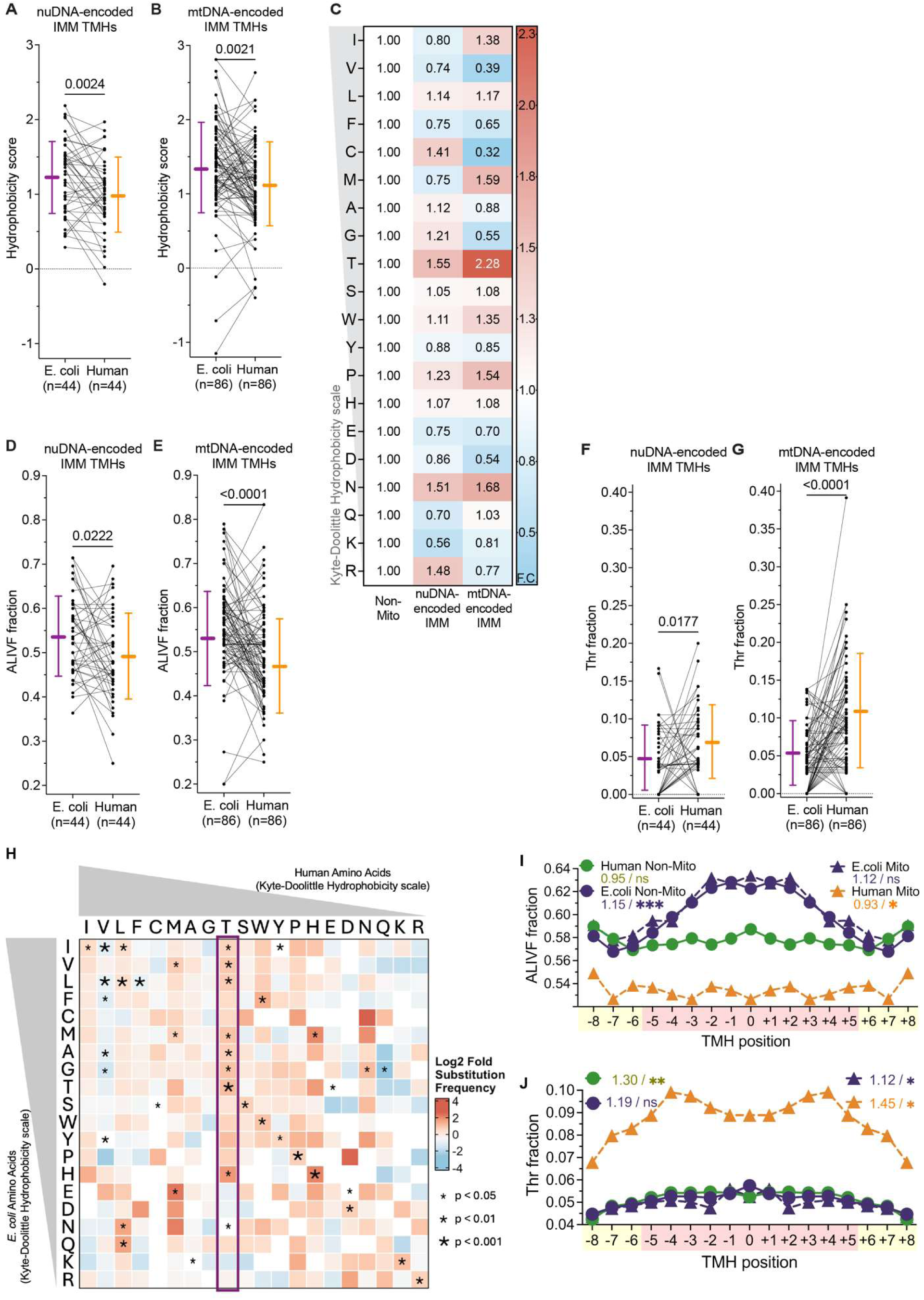
Evolutionary comparisons show aliphatic-to-threonine substitutions in mitochondrial transmembrane proteins. **(A, B)** Hydrophobicity scores of structurally aligned TMH pairs from evolutionarily related *E. coli* and *H. sapiens* nuDNA-encoded and mtDNA-encoded IMM proteins, respectively. **(C)** Mean amino acid composition of TMHs from evolutionarily related *H. sapiens* nuDNA-encoded and mtDNA-encoded IMM proteins, expressed as fold change relative to paired *E. coli* TMHs and normalized to the corresponding fold change in non-mitochondrial TM proteins. Red indicates enrichment, blue indicates depletion. **(D, E)** Aliphatic residue fractions of structurally aligned TMH pairs from evolutionarily related *E. coli* and *H. sapiens* nuDNA-encoded and mtDNA-encoded IMM proteins, respectively. **(F, G)** Threonine fractions of structurally aligned TMH pairs from evolutionarily related *E. coli* and *H. sapiens* nuDNA-encoded and mtDNA-encoded IMM proteins, respectively. Lines connect paired helices; bars represent mean ± SD. p-values from Wilcoxon signed-rank test. **(H)** Relative substitution matrix for aligned TMHs from evolutionarily related *E. coli* and *H. sapiens* IMM TM proteins, normalized to substitutions from non-IMM TMHs. Asterisks denote statistical significance from Fisher’s exact test. **(I, J)** Positional distribution of ALIVF fraction (I) and threonine fraction (J) along UniProtKB-annotated TMHs from *H. sapiens* mitochondrial and non-mitochondrial TM proteins and their respective evolutionarily related *E. coli* proteins, aligned agnostically to membrane orientation. Values on figure denote odds ratio/significance for core vs. peripheral enrichment from Fisher’s exact test (✱p<0.05, ✱✱p<0.01, ✱✱✱p<0.001).

The above pairwise comparisons, by definition, only analyzed proteins that were evolutionarily related. To test generalizability across the entire human IMM and *E. coli* proteomes, we performed similar analyses across all UniProtKB-annotated human and *E. coli* transmembrane proteins. These proteome-wide analyses recapitulated all key trends observed in pairwise comparisons: reduced TMH hydrophobicity, decreased aliphatic residue fraction, and increased threonine content in human IMM proteins (Figure 2—figure supplement 3A–C). We applied this proteome-level approach to compare the human IMM proteome with that of *R. rickettsii*, a relative of the ancestral mitochondrial endosymbiont (Kurland and Andersson, 2000). Comparison of human IMM proteins with *R. rickettsii* showed similar results (Figure 2—figure supplement 3D–F), consistent with these compositional adaptations being a robust and conserved feature of mitochondrial protein evolution. To test if ATS is observed in other endosymbiotic organelles, we performed a similar analysis on *A. thaliana* chloroplast thylakoid proteins, which face analogous cytosolic solubility constraints to those of human IMM proteins. As a comparator, we used the cyanobacterium *Synechocystis*, which is related to the ancestral chloroplast (Sato, 2020). While thylakoid TMHs also showed a modestly reduced hydrophobicity, threonine enrichment was not observed (Figure 2—figure supplement 3G–I). This could reflect less severe membrane insertion constraints, because chloroplasts retain a Sec-dependent biogenesis machinery that is functionally similar to that of the endoplasmic reticulum and enables co-translational protein insertion (Ballabani et al., 2023).

As human inner mitochondrial membrane TMHs showed reduced aliphatic and increased threonine content, we next queried whether this is the result of specific amino acid substitutions. We generated a relative substitution matrix comparing mitochondrial against non-mitochondrial TMHs (Figure 2H). Threonine ranked among the most conserved residues — second only to histidine, and ahead of proline and tryptophan, residues typically constrained by their unique structure. Our analyses revealed significant enrichment of aliphatic-to-threonine substitutions (ATS) in mitochondrial TMHs. This signal was TMH-specific, since similar analyses for soluble helices from either the same TM proteins or from fully soluble proteins did not show similar ATS enrichment (Figure 2—figure supplement 2A, B). Further, threonines were enriched, and aliphatic residues depleted, in the hydrophobic core of the TMH (Figure 2I, J) — a distribution consistent with ATS-driven hydrophobicity reduction. Together, these *E. coli*-human protein comparisons support a role of ATS in decreasing hydrophobicity in this set of evolutionarily conserved human IMM proteins, raising the question of whether solubility-stability constraints are also evident across the broader human proteome.

### Mitochondrial transmembrane proteins have a distinct amino acid composition

We next compared hydrophobicity and residue composition of structure-derived TMHs across human transmembrane proteins of a shared nuclear origin but different cellular destinations (Figure 1A, Figure 3A). Non-mitochondrial TMHs were remarkably uniform: endoplasmic reticulum, Golgi, cell membrane, and nuclear envelope proteins showed similar TMH properties despite their distinct lipid and biochemical environments. Mitochondrial TMHs stood apart, however, with lower hydrophobicity, lower aliphatic fraction, and higher threonine content (Figure 3A–D, Figure 3—figure supplement 1A–D). To ensure a like-for-like comparison, beta-barrel proteins were excluded from the analysis. We saw the same differences when we analyzed all human transmembrane proteins using UniProtKB-defined TMH annotation (Figure 3—figure supplement 2A–D).

**Figure 3:**
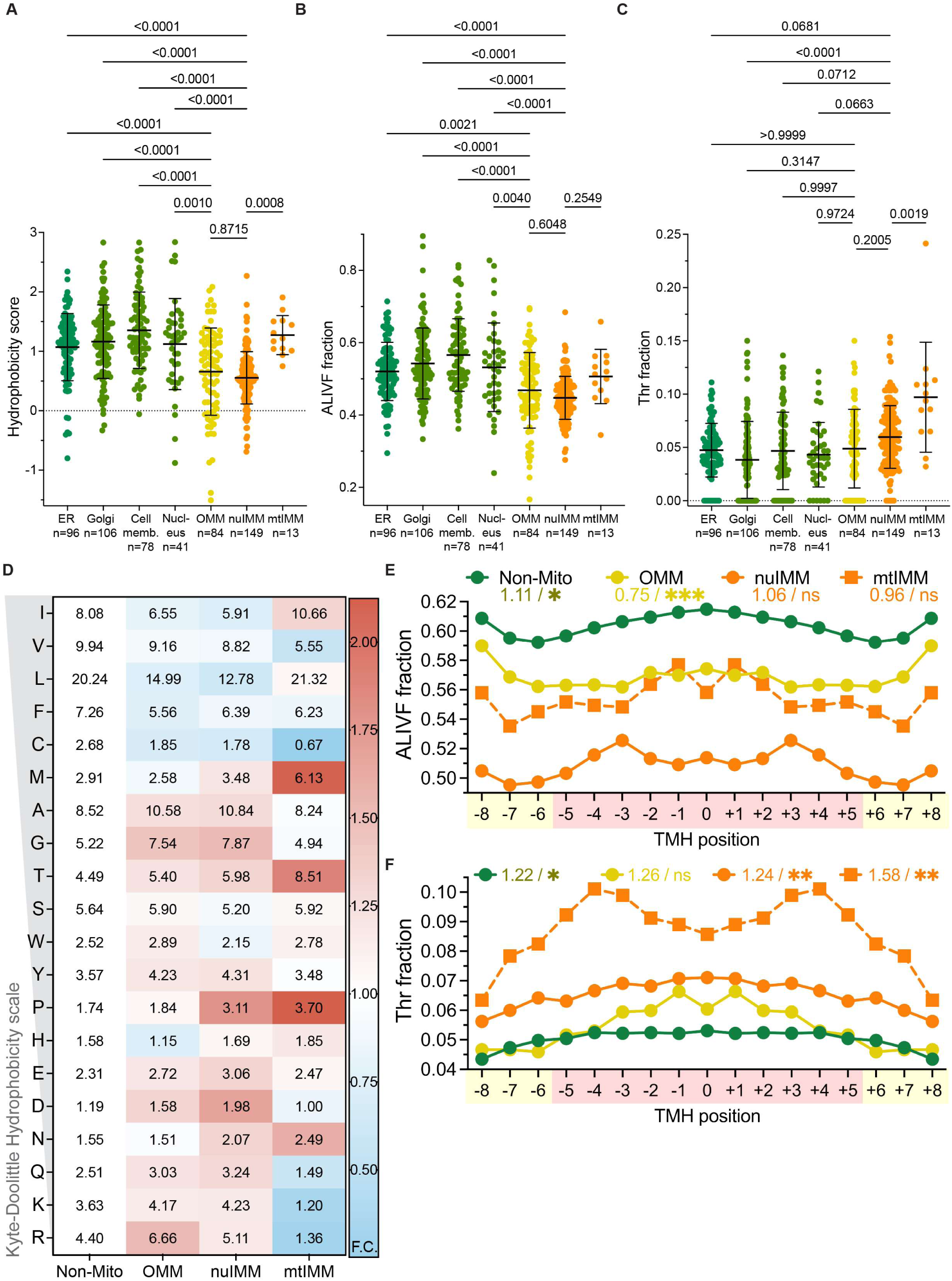
Threonine enrichment distinguishes human mitochondrial proteins. **(A–C)** Hydrophobicity scores (A), ALIVF fraction (B), and threonine fraction (C) of human TMHs identified from TM protein AlphaFold2 structures, grouped by membrane compartment. Each point represents one TM protein; bars represent mean ± SD. p-values from one-way ANOVA with Tukey’s post-hoc test (all pairwise comparisons shown in Figure 3—figure supplement 1). **(D)** Amino acid residue frequency (%) in human TMHs identified from TM protein AlphaFold structures, grouped by membrane compartment. Cell values represent mean residue frequency; color scale reflects fold-change relative to the mean of non-mitochondrial compartments (red, enriched; blue, depleted; all compartments shown separately in Figure 3—figure supplement 1). **(E, F)** Positional distribution of ALIVF fraction (E) and threonine fraction (F) along TMHs from human TM proteins, aligned agnostically to membrane orientation, grouped by membrane compartment. Values on figure denote odds ratio/significance for core vs. peripheral enrichment from Fisher’s exact test (✱p<0.05, ✱✱p<0.01, ✱✱✱p<0.001).

We examined residue positions along the length of TMHs and found that aliphatic residues were enriched in the helix core of non-mitochondrial TMHs but not in mitochondrial ones, which were instead enriched for threonine (Figure 3E, F). Structure-based orientation of TMHs gave the same core pattern of aliphatic depletion and threonine enrichment (Figure 3—figure supplement 3B, C). Hydrophilic residues, by contrast, were enriched in peripheral regions (Figure 3—figure supplement 3A), consistent with threonine rather than other hydrophilic residues driving TMH hydrophobicity reduction.

Several independent controls confirmed the specificity of threonine enrichment to mitochondrial transmembrane helices. Soluble helices showed no compositional differences across compartments, whether from the same TM proteins or from fully soluble proteins (Figure 3— figure supplement 4A–D, 5A–D). Comparison within individual mitochondrial proteins, an approach that controls for all protein-specific confounders, showed threonine was enriched in transmembrane over soluble helices (Figure 3—figure supplement 6A–H). The enrichment was threonine-specific and not a general enrichment of polar residues. For example, serine, which is most chemically similar to threonine, was enriched across all non-mitochondrial helices and even in mitochondrial soluble helices, but in mitochondrial transmembrane helices threonine was favored over serine (Figure 3—figure supplement 7).

We next expanded the scope of our analyses to other eukaryotic groups. Across metazoans, where mitochondrial targeting challenges are shared with humans, mitochondrial transmembrane proteins likewise displayed ATS (Figure 3—figure supplement 8A, D). In higher plants, which possess two endosymbiosis-derived organelles and therefore face a solubility-stability tradeoff for both mitochondrial and chloroplast thylakoid membrane proteins, TMHs showed reduced hydrophobicity without evidence of marked threonine enrichment in either organelle (Figure 3—figure supplement 8B, E). A similar pattern was observed in jakobids which did not show increased mitochondrial TMH threonine content despite reduced hydrophobicity (Figure 3—figure supplement 8C, F). This is likely due to the atypical retention of additional proteobacteria-derived insertion machineries in plants and jakobids (Ballabani et al., 2023; Kolli et al., 2020; Moreira et al., 2024). Collectively, these findings indicate that ATS is specific to transmembrane mitochondrial proteins, with their unique challenges in solubility and stability.

### ATS accompanies nuclear relocation of mitochondrial genes across eukaryotes

To investigate the relationship between ATS and mitochondrial targeting constraints more broadly throughout the eukaryotic kingdom, we leveraged examples of evolutionary gene transfer from mtDNA to the nuclear genome (Butenko et al., 2024) (Figure 4A). For each mtDNA-encoded protein, we identified two closely related eukaryotic organisms: one retaining the mtDNA-encoded version, and another in which the same protein is nuDNA-encoded (Figure 4B). These matched pairs were selected to maximize evolutionary proximity while allowing direct comparison of functional mtDNA- and nuDNA-encoded protein variants in extant lineages. All protein pairs were then structurally aligned to identify compositional changes.

**Figure 4:**
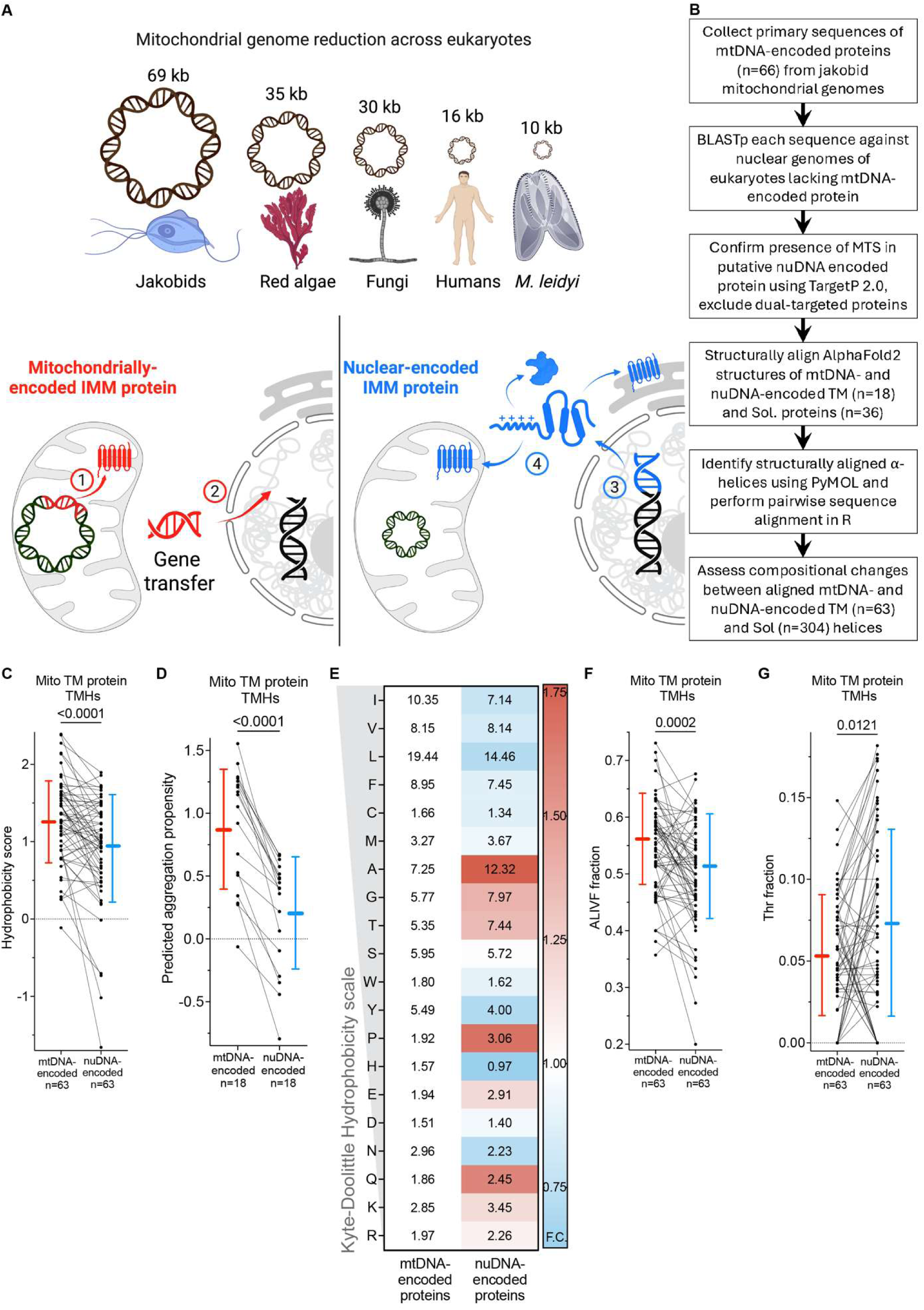
ATS is found in mitochondrial gene transfers. **(A)** Top, mtDNA reduction across eukaryotes reflects differential retention of mitochondrial genes after endosymbiotic gene transfer. Bottom, mtDNA-encoded IMM proteins are synthesized and inserted locally in the matrix (1). After endosymbiotic gene transfer to the nuclear genome (2), the same IMM protein becomes nuDNA-encoded (3) and requires mitochondrial targeting (4), exposing hydrophobic TMHs to aggregation and mistargeting. **(B)** Computational pipeline for identifying and analyzing mtDNA-to-nuDNA gene transfer events across eukaryotic phyla. **(C, D)** Hydrophobicity scores (C) and predicted aggregation propensity (D) of structurally aligned TMH pairs from mtDNA- and nuDNA-encoded versions of the same IMM protein across eukaryotes. **(E)** Amino acid residue frequency (%) in TMHs of mtDNA- and nuDNA-encoded IMM proteins across eukaryotes. Cell values represent mean residue frequency; color scale reflects fold-change relative to mtDNA-encoded proteins (red, enriched; blue, depleted). **(F)** ALIVF fraction and **(G)** threonine fraction of structurally aligned TMH pairs from mtDNA- and nuDNA-encoded versions of the same IMM protein across eukaryotes. Lines connect paired helices; bars represent mean ± SD. p-values from Wilcoxon signed-rank test.

For nuDNA-encoded proteins, transmembrane helices exhibited reduced hydrophobicity, whereas soluble helices from the same proteins or from soluble proteins showed no significant change (Figure 4C, Figure 4—figure supplement 1A, B). Because efficient mitochondrial targeting also depends on aqueous compatibility, we performed a structure-based aggregation analysis. All nuclear genome-transferred transmembrane proteins showed reduced predicted aggregation propensity, whereas relocated soluble proteins showed no reduction and instead exhibited a modest increase (Figure 4D, E, Figure 4—figure supplement 1C, D). We next examined compositional changes underlying the reduced hydrophobicity and predicted aggregation propensity during gene transfer. Aliphatic residue fraction was reduced in transmembrane helices but remained unchanged in soluble helices from both transmembrane and soluble proteins (Figure 4F, Figure 4—figure supplement 1E, F). This was accompanied by an increased threonine fraction in TMHs, whereas soluble helices from both protein classes showed no change, consistent with the ATS we observed in previous analyses (Figure 4G, Figure 4—figure supplement 1G, H). To control for genome-wide composition bias across our phylogenetically diverse organismal pairs, we calculated the ratio of mean TMH threonine content in nuDNA-encoded to mtDNA-encoded TM proteins for each organism and found that it did not differ significantly (Figure 4—figure supplement 1I). Positional analysis along TMHs further showed that aliphatic depletion and threonine enrichment were both concentrated in the core regions of nuclear-encoded TMHs relative to their mtDNA-encoded counterparts (Figure 4—figure supplement 1J, K). The gene-transfer pairs showing ATS include green algae. As with metazoans, green algae retain minimal insertase machinery, unlike the expanded Oxa family found in land plants (Figueroa-Martínez et al., 2008). This more robust insertion machinery is consistent with land-plant mitochondrial TMHs not necessitating threonine enrichment despite having shared mitochondrial ancestry with green algae that possess reduced IMM protein biogenesis machinery (Figure 3—figure supplement 8B, E).

We next asked whether ATS is evident at the level of individual amino acid substitutions in gene transfers from the mtDNA to the nuclear genome. A standard substitution matrix analysis was not feasible because gene transfers in the reverse direction (nuclear to mtDNA) are not observed in evolution, leaving no appropriate control (Butenko et al., 2024). We therefore analyzed ALIVF⇔X interconversion frequencies in TMHs, which should be comparable in the absence of any directional evolutionary pressure. Aliphatic-to-threonine substitutions were favored over the reverse direction during transmembrane protein-encoding gene transfers (OR = 2.13; P_adj._ = 6.84 × 10⁻¹⁰), consistent with ATS. Together, these gene-transfer analyses show that transmembrane proteins undergo ATS upon nuclear relocation, most likely to facilitate mitochondrial targeting.

### Convergent ATS acquisition in distinct ATP6 gene transfer events

Our gene transfer analysis identified several instances in which the same gene became nuclear encoded in multiple distantly related eukaryotic lineages. *MT-ATP6*, which encodes a protein of the ATP synthase proton channel, is absent from the mtDNA of apicomplexans, chlorophytes and metazoan ctenophores, and is instead nuclear encoded in these species (Funes et al., 2002; Mühleip et al., 2021; Pett et al., 2011). Phylogenetic analysis showed that the ATP6 sequences from these three clades did not segregate into a single monophyletic group (Figure 5A). Instead, nuclear-encoded sequences grouped with mitochondrially encoded ATP6 sequences from the same clade, forming three distinct, well-supported monophyletic groups. TargetP 2.0 MTS analysis (Almagro Armenteros et al., 2019) further showed that nuclear-encoded ATP6s in each clade carried highly divergent MTSs with differing predicted cleavage sites (Figure 5—figure supplement 1A–C). Together, these data strongly suggest three independent transitions to the nuclear genome. Similar phylogenetic and MTS analyses of another mitochondrially-encoded gene, *MT-ATP9,* supported four independent gene transfers in apicomplexans, metazoan ctenophores, fungi, and chlorophytes (Figure 5—figure supplement 2, 3A–D).

**Figure 5:**
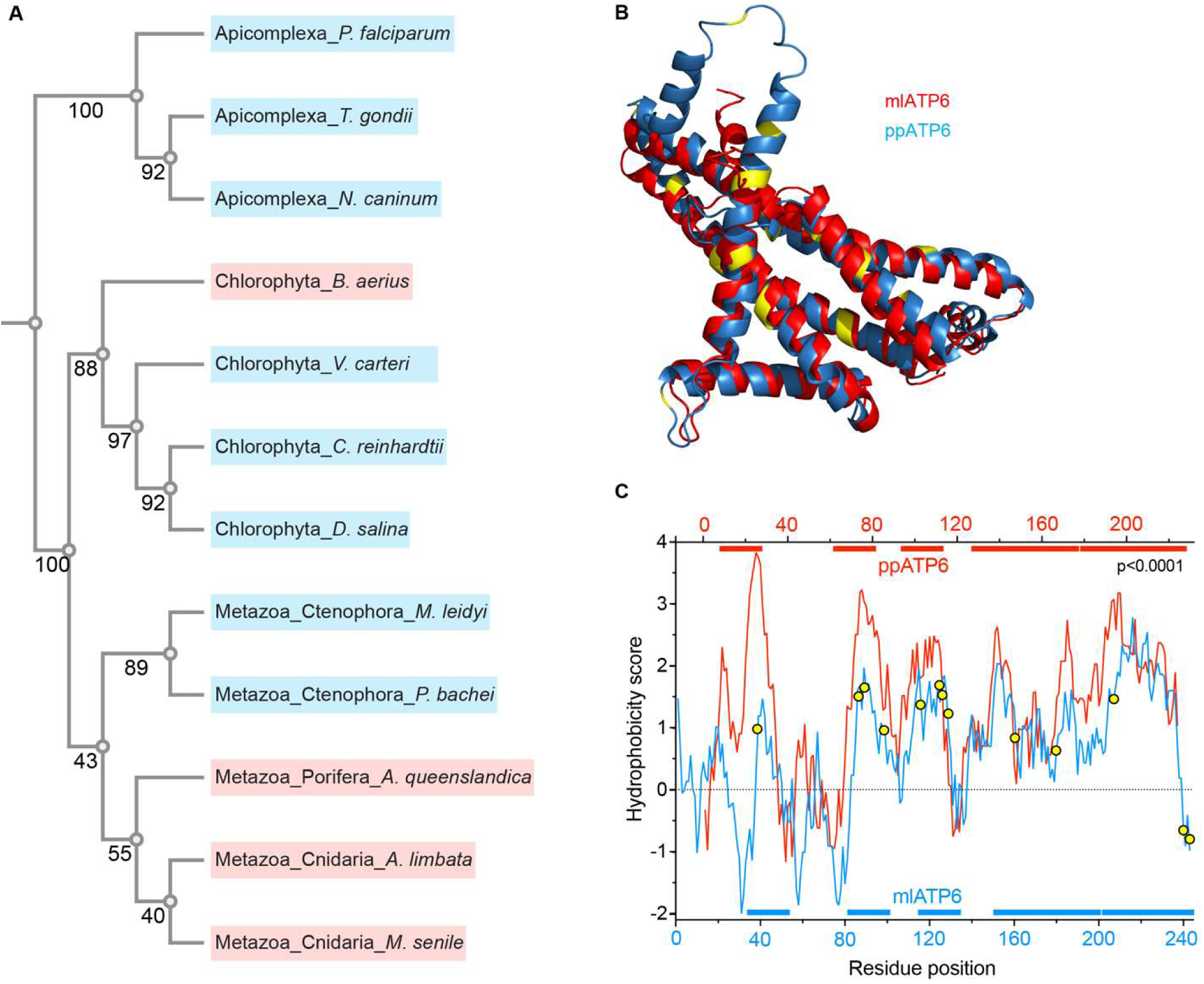
ATS is convergently acquired in independent ATP6 gene transfer events. **(A)** Phylogenetic tree of ATP6 sequences from apicomplexans, chlorophytes, and metazoans. Blue indicates nuclear-encoded ATP6; red indicates mitochondrially encoded ATP6. Numbers at nodes indicate bootstrap support values. **(B)** Structural alignment of mlATP6 (blue) and ppATP6 (red) reveals conservation of proton channel architecture following nuclear gene transfer. Yellow indicates threonine residues in mlATP6. RMSD = 1.790 Å. **(C)** Whole-protein hydrophobicity profiles of mlATP6 (blue) and ppATP6 (red). Colored bars above and below indicate predicted transmembrane helices for ppATP6 and mlATP6, respectively. Yellow circles indicate threonine residues in mlATP6 TMHs. p<0.0001, Wilcoxon rank-sum test comparing TMH hydrophobicity scores between mlATP6 and ppATP6.

To test whether convergent compositional adaptations are found in these diverse clades, we compared the nuclear-encoded ATP6 from the ctenophore *Mnemiopsis leidyi* (ml) with a mitochondrially-encoded ATP6 from the closely related cnidarian *Porites panamensis* (pp). Structural alignment of mlATP6 and ppATP6 revealed strong conservation of the proton channel architecture (Figure 5B). Because mtDNA-encoded ATP6 is a hydrophobic TM protein, we next assessed whether its nuclear transfer was accompanied by reduced TMH hydrophobicity through ATS. Indeed, mlATP6 exhibited reduced TMH hydrophobicity relative to ppATP6, accompanied by threonine enrichment in TMH regions (Figure 5C). Pairwise sequence alignments confirmed threonine substitutions in mlATP6 at multiple aliphatic positions of ppATP6, consistent with ATS (Figure 5—figure supplement 4A). Examination of the two additional examples of ATP6 nuclear genome translocation in apicomplexans and chlorophytes also showed reduced TMH hydrophobicity, Thr enrichment and clear evidence of ATS (Figure 5—figure supplement 4B–G).

### ATS is required for mlATP6 mitochondrial targeting

Having identified ATS in multiple independent ATP6 mtDNA-to-nuDNA transfer events, we next tested whether mlATP6 targets efficiently to mitochondria when expressed in mammalian cells. We expressed mlATP6 with a variety of C-terminal tags (EGFP, V5, Neptune) in multiple mammalian cell lines (HEK293T, Neuro2A, C2C12, JHH7) and observed robust mitochondrial localization across all constructs and cell lines tested (Figure 6—figure supplement 1A–E). To test whether mlATP6 targeting requires ATS, we expressed three ATP6 constructs: 1) WT-hATP6 (human MT-ATP6 with the mlATP6 MTS appended) with high native hydrophobicity; 2) WT-mlATP6, with native ATS and low hydrophobicity; and 3) STA-mlATP6 carrying Substitutions of Threonine to Aliphatic residues (STA) that reverse ATS and thereby increase hydrophobicity (Figure 6A, 6B). In HEK293T cells, WT-hATP6 was extensively mislocalized despite carrying the mlATP6 MTS, whereas WT-mlATP6 was robustly targeted to mitochondria (Figure 6C, D). Simply reversing ATS in the STA-mlATP6 protein was sufficient to cause its mislocalization (Figure 6E–G, Figure 6—figure supplement 1F). We therefore conclude that ATS is critical for efficient mitochondrial targeting in a native example of ATP6 nuclear gene transfer.

**Figure 6:**
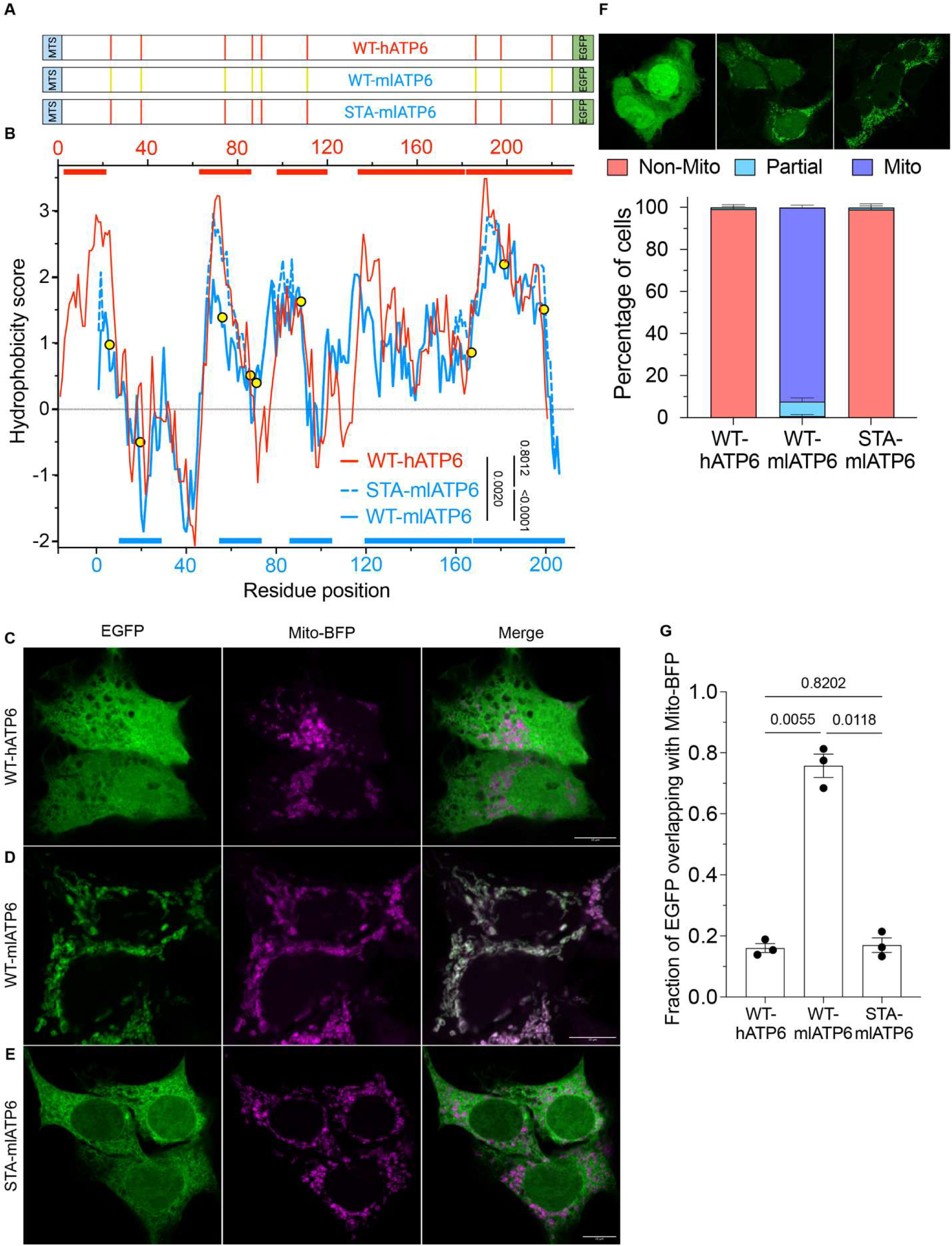
ATS is required for mlATP6 targeting. **(A)** Schematic of the three EGFP-tagged ATP6 constructs: WT-hATP6 (human MT-ATP6 with mlATP6 MTS appended), WT-mlATP6, and STA-mlATP6 (Thr-to-aliphatic substitutions reversing ATS). Yellow lines, threonine residues in WT-mlATP6; red lines, Thr-to-aliphatic substitutions in STA-mlATP6. **(B)** Whole-protein hydrophobicity profiles of WT-hATP6 (red), WT-mlATP6 (solid blue), and STA-mlATP6 (dashed blue). Yellow circles mark threonine positions in WT-mlATP6. Bars above and below indicate TMH regions; p-values denote hydrophobicity comparisons between constructs. **(C–E)** Confocal imaging of HEK293T cells expressing WT-hATP6 (C), WT-mlATP6 (D), or STA-mlATP6 (E) shown as EGFP (green), Mito-BFP (magenta), and merged channel. Scale bars, 10 µm. (**F**) Scoring of EGFP localization across cells as Non-Mito, Partial, or Mito for each construct. Data are mean ± SD from three independent experiments. Representative images of each scoring category (Non-Mito, Partial, Mito; left to right) are shown above. (**G**) Manders’ colocalization coefficient (M1; fraction of EGFP overlapping with Mito-BFP) quantifying mitochondrial localization for each construct. Data are mean ± SD from three independent experiments; p-values from repeated-measures one-way ANOVA with Greenhouse-Geisser correction and Tukey’s multiple comparisons test.

### ATS enables mitochondrial targeting of nuclear-encoded hATP6

Given that expression of mtDNA-encoded proteins from the nuclear genome has proven very challenging, we next evaluated whether ATS could be harnessed to enable mitochondrial targeting of hydrophobic IMM proteins. We selected the mtDNA-encoded hATP6, given its high hydrophobicity and mislocalization when expressed from the nuclear genome in human cells, as observed both in our experiments and in published studies (Artika, 2020; Björkholm et al., 2017). We first asked whether previously employed strategies such as codon optimization or appending strong non-mammalian MTSs — which do not directly lower protein hydrophobicity — would be sufficient to enable hATP6 targeting. We appended a variety of N-terminal MTSs to codon-optimized hATP6-WT but observed limited mitochondrial targeting in HEK293T cells (Figure 7— figure supplement 1).

Therefore, we leveraged ATS to lower TMH hydrophobicity, designing a series of hATP6 constructs with progressively increasing aliphatic-to-threonine substitutions (Figure 7A). To minimize structural perturbations, ATS swaps were restricted to aliphatic positions where threonine is naturally present in at least one ATP6 sequence in the multiple sequence alignment. AlphaFold models of the ATP6 structures are shown in Figure 7—figure supplement 2A, B. The resulting hATP6-ATS constructs (T1, T2, T3) showed progressively reduced protein hydrophobicity and reduced predicted aggregation propensity with increasing threonine content (Figure 7B, Figure 7—figure supplement 2C).

**Figure 7:**
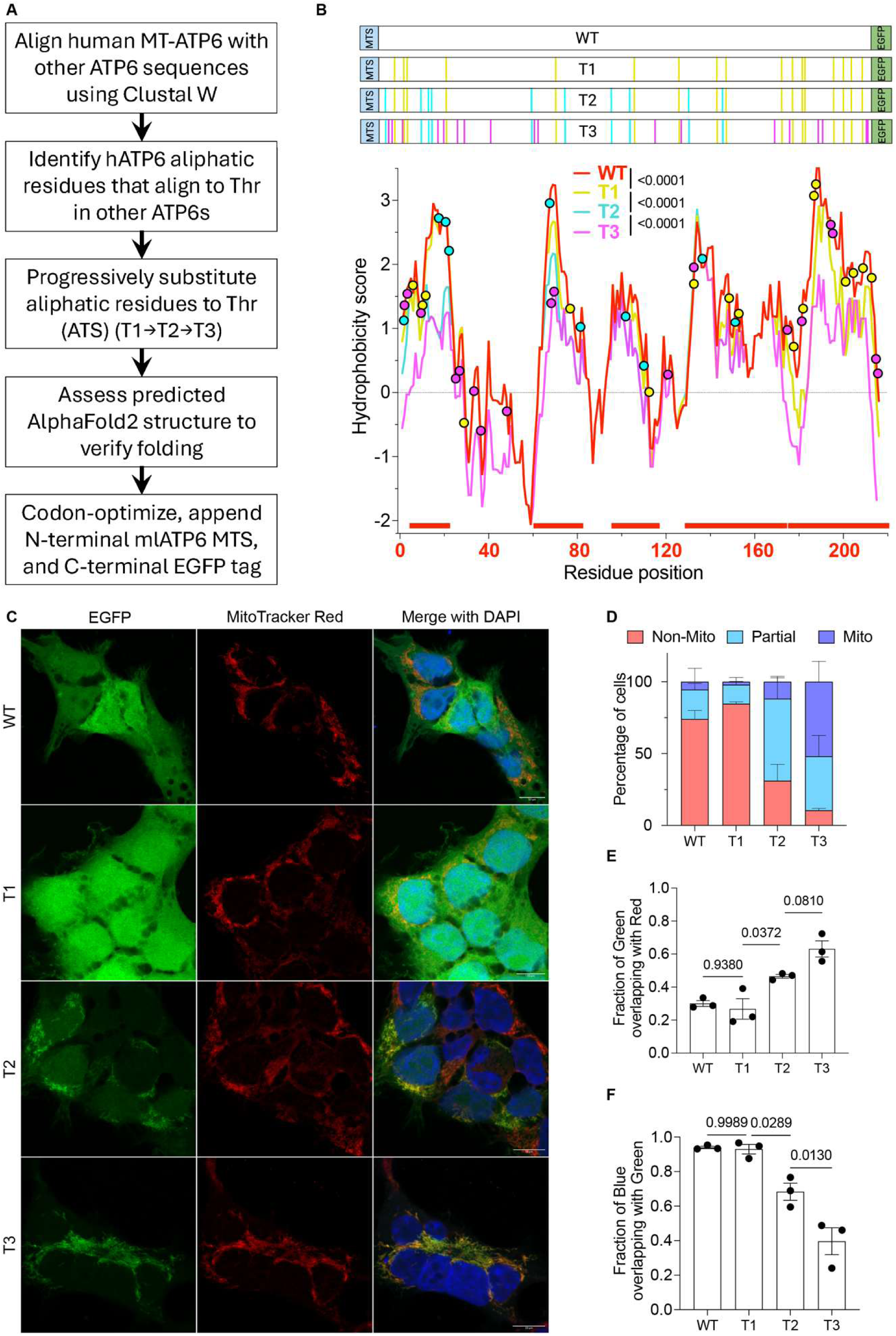
ATS supports hATP6 targeting. **(A)** Computational pipeline for designing progressively less hydrophobic hATP6 constructs. **(B)** Schematics of the hATP6-WT, -T1, -T2, and -T3 constructs (MTS, blue; ATP6, white; EGFP, green); vertical lines mark aliphatic-to-threonine substitution positions, colored by construct (T1, yellow; T2, cyan; T3, magenta). Below, Kyte-Doolittle hydrophobicity profiles of each construct with threonine positions indicated. p-values from pairwise Wilcoxon rank-sum tests comparing sequential constructs. **(C)** Representative confocal images of HEK293T cells stably expressing the indicated hATP6 constructs (EGFP, green; MitoTracker Red, red; nuclei, blue), shown as individual and merged channels. Scale bars, 10 µm. **(D)** Scoring of EGFP localization across cells as Non-Mito, Partial, or Mito for each construct. **(E)** Manders’ colocalization coefficient (M1; fraction of EGFP overlapping with MitoTracker Red) for each construct. **(F)** Fraction of DAPI (nuclear) signal overlapping with EGFP for each construct, reflecting non-mitochondrial EGFP distribution. Data are mean ± SEM from three independent experiments; p-values from one-way ANOVA with Tukey’s post hoc test.

Upon stable expression in HEK293Ts, hATP6-ATS constructs showed improved mitochondrial targeting with increasing ATS implementation, as quantified by colocalization analyses (Figure 7C–F). Additionally, we controlled this comparison by biasing it against our hypothesis. From a polyclonal population of cells expressing hATP6-ATS, we selected lower-expressing cells for WT and T1 and higher-expressing cells for T2 and T3. Despite expressing at higher levels, T2 and T3 showed higher fractional mitochondrial targeting, whereas WT and T1 remained mislocalized even at lower expression levels (Figure 7—figure supplement 3). Mitochondrial targeting is therefore driven primarily by hydrophobicity rather than simply being a function of low expression. Together, these results show that introducing threonine residues in place of aliphatic residues, when combined with an appropriate MTS, is sufficient to enable mitochondrial targeting of a highly hydrophobic mtDNA-encoded IMM protein when expressed from the nuclear genome.

### Threonine remains membrane-compatible despite its polar character

To understand how threonine remains compatible with the membrane interior despite its polar hydroxyl moiety, we used MD simulations of two TM proteins of the IMM to examine threonine sidechain rotamer behavior across different solvation environments. We analyzed ATP/ADP carrier protein 1 (ANT1) and uncoupling protein 1 (UCP1), each simulated over six independent runs (runtime ∼6 μs) in a 1-palmitoyl-2-oleoyl-sn-glycero-3-phosphocholine (POPC) bilayer containing 3 protein-associated cardiolipins and 7 mol% arachidonic acid (Figure 8A). We analyzed threonine conformation in three combinations of structural and solvent environments: alpha helices facing the membrane core (as would occur in a folded TMH of an IMM protein), loops facing water (a model for the unfolded protein in transit to the mitochondria), and alpha helices facing water (to separate the effects of solvent environment and secondary structure).

**Figure 8:**
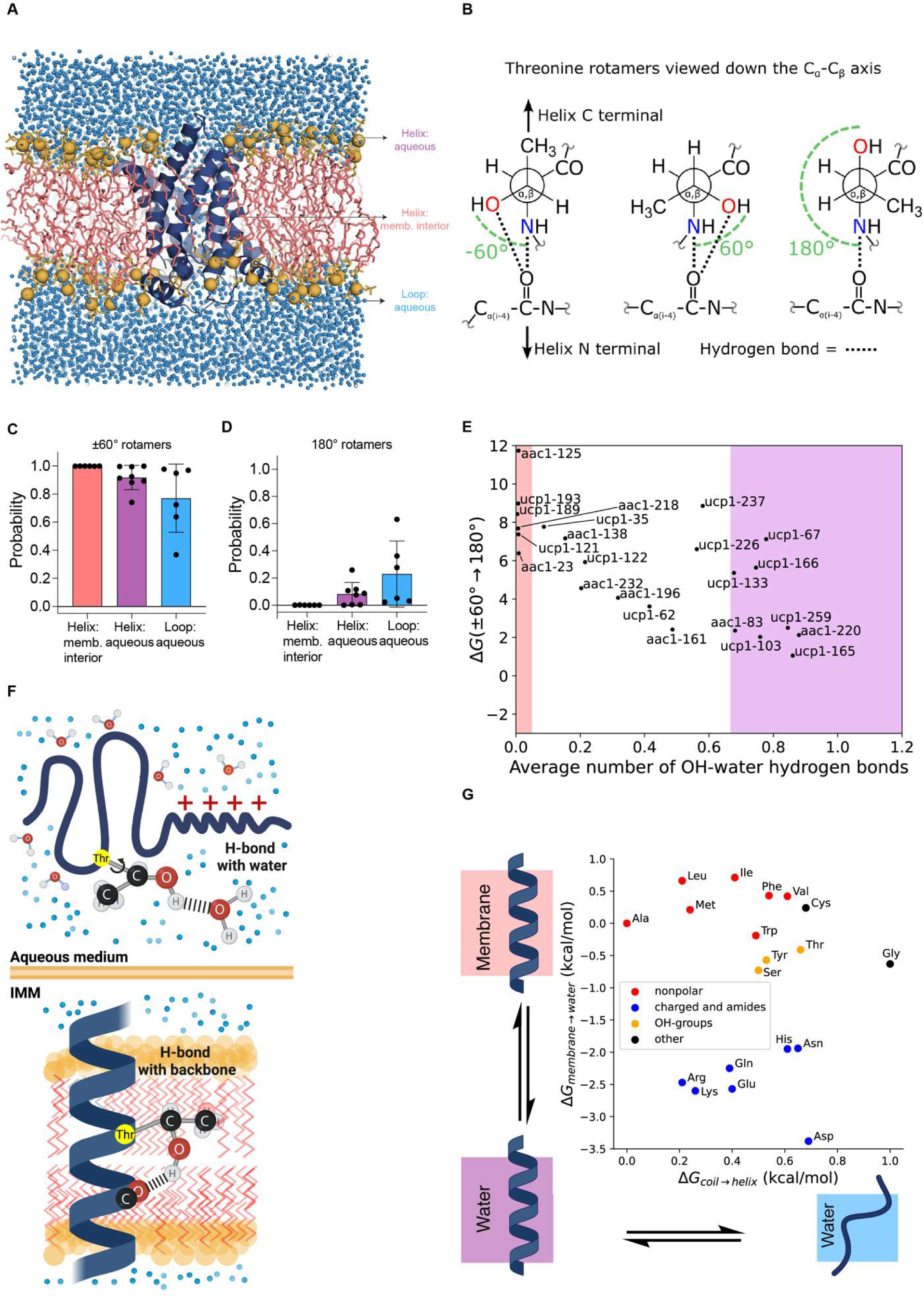
Threonine remains membrane-compatible despite polar character. **(A)** Schematic of simulation system, with UCP1 in dark blue ribbons, and POPC membrane in red sticks. The figure was made using PyMOL 3.1.6.1. **(B)** Canonical χ1 rotamers of threonine. Newman projections along the Cα–Cβ bond. Red, Oγ; blue, backbone N; dashed arc, χ1 dihedral. At −60° and +60°, the hydroxyl oxygen is positioned to form a hydrogen bond with the backbone carbonyl; the 180° rotamer precludes this interaction. **(C, D)** χ1 rotamer occupancy of threonine residues by structural context. Mean probability (±SD) of occupying the ±60° or 180° χ1 rotamer for threonines in UCP1 and ANT1, grouped by context: membrane-interior helix (red, n=6), aqueous helix (blue, n=8), and aqueous loop (cyan, n=6). Dots represent individual residues. **(E)** Rotamer transition free energy as a function of OH–water hydrogen bonding. ΔG(±60°→180°) for individual threonine residues in UCP1 and ANT1 plotted against their mean number of hydroxyl–water hydrogen bonds. Red and blue shading denote the membrane-interior and aqueous environments, respectively. **(F)** Model for threonine’s role in ATS: its hydroxyl hydrogen bonds to water while the protein is unfolded during cytoplasmic transport, and to the i−4 backbone carbonyl once folded into transmembrane helices — keeping threonine compatible with both mitochondrial targeting and transmembrane helix folding. **(G)** Membrane-to-water transfer free energy versus helical propensity for all amino acids. ΔG(membrane → water) plotted against ΔG(coil → helix) for each amino acid, colored by side-chain chemistry: nonpolar (red), charged and amides (blue), OH-groups (orange), and other (black). Schematics depict the membrane-to-water and coil-to-helix equilibria. Proline is absent in the published helicity scale due to its extremely unfavorable free energy of helix formation and was omitted from this figure.

In an α-helix, the −60° and +60° rotamers of the threonine Cα–Cβ bond position the sidechain hydroxyl to hydrogen bond with the backbone carbonyl at position i−4, whereas in the 180° rotamer the hydroxyl faces outward and is more exposed to solvent (Figure 8B). We therefore hypothesized that the ±60° rotamers would be favored in membrane-interior TM helices, while the 180° rotamer, which cannot form this backbone hydrogen bond, would be disfavored there. In helical regions facing the membrane interior, threonine strongly favored the ±60° rotamers, whereas the 180° rotamer was essentially unpopulated (Figure 8C, D), reflecting the energetic penalty associated with exposing the sidechain hydroxyl to the hydrophobic fatty acid tails of membrane phospholipids. In aqueous helical regions, threonine adopted the 180° rotamer configuration to some extent, but still mostly populated the ±60° rotamers. In unstructured loop regions, which have the greatest aqueous exposure, the relative probabilities of the ±60° and 180° rotamers were statistically indistinguishable (from the 1:1:1 ratio of −60°, +60°, and 180° rotamers expected if all were equally probable), consistent with the flexibility of loops.

We next analyzed the free energy cost of transitioning from ±60° to the 180° rotamer, ΔG(±60°→180°), estimated for each residue from the fraction of simulation frames spent in each rotameric state (see Methods). We grouped threonine into three solvation categories — membrane-interior, aqueous-exposed, and intermediate — based on the average number of hydrogen bonds between the sidechain hydroxyl and water (Figure 8E). In general, membrane-interior threonines had higher ΔG(±60°→180°) than aqueous-exposed threonines (Figure 8E), consistent with the destabilization expected if the sidechain hydroxyl were exposed to the membrane interior. Averaging across residues in the membrane and aqueous categories, the free energy of the 180° rotamer is expected to increase by ∼4.9 kT (95% CI = 2.5–7.3 kT) upon membrane insertion.

To independently estimate the effect of the membrane environment on threonine rotamers, we constructed a thermodynamic cycle from published threonine sidechain-analogue and whole-residue membrane insertion free energies (details in Methods). The resulting change in the free energy of rotation from the ±60° rotamers to the 180° rotamer was ∼4.5 kT, similar to the estimate of ∼4.9 kT from MD simulations.

Together, these results support a model (Figure 8F) in which the hydroxyl group allows threonine to form energetically favorable hydrogen bonds in both aqueous and membrane environments, which IMM proteins encounter in their life cycle. During post-translational transit through the cytoplasm, the hydroxyl can hydrogen bond with surrounding water. As the helix is inserted into the membrane interior, where water is largely excluded, threonine almost exclusively adopts the ±60° rotamer and hydrogen bonds with the i−4 backbone carbonyl, thereby avoiding the energetic cost of exposing an unsatisfied polar hydroxyl to the hydrophobic membrane interior. We propose that its intermediate hydrophobicity and low (but not extreme like that of glycine and proline) helical propensity (Figure 8G) combine to make threonine uniquely compatible with solubility in the cytoplasm, transit through the mitochondrial import machinery, and helix folding in the mitochondrial inner membrane.

## DISCUSSION

Mitochondrial membrane proteins face a unique biophysical challenge: combining aqueous solubility with stability within the hydrophobic membrane. We report aliphatic-to-threonine substitution as a recurrent residue-level adaptation of inner mitochondrial membrane proteins that is likely a solution to this challenge. Solving this tradeoff involves satisfying conflicting constraints: lower hydrophobicity can reduce aggregation risk during targeting, but strongly polar or charged substitutions would make membrane insertion less favorable (Hessa et al., 2005; White and Wimley, 1999). Threonine fits this constraint because it is polar enough to lower hydrophobicity without introducing a charged sidechain that would compromise membrane compatibility. This interpretation is consistent with the observation that mtDNA-encoded IMM proteins retain higher TMH hydrophobicity than nuclear-encoded IMM proteins but still gain threonine while avoiding the most hydrophilic residues. The spatial pattern of residues in the membrane reinforces this. Threonine is most strongly enriched in core regions where sidechains typically face either nonpolar lipid tails or other hydrophobic protein segments, whereas other polar residues are enriched at peripheral positions where charged and amide-bearing sidechains can rotate to reach the more polar interfacial region. At the atomic level, our MD analyses explain how threonine can support this dual character. It can lower hydrophobicity in the aqueous environment by hydrogen bonding with solvent, while offsetting the energetic cost of burying a polar group through backbone hydrogen bonding.

The examples of *MT-ATP6* gene transfer to the nuclear genome provide a natural comparative test of this model. In apicomplexans, chlorophytes, and ctenophores, independently nuclear-transferred *MT-ATP6* genes all show ATS. The contrast between divergent MTSs and convergent ATS is notable: targeting peptides were acquired independently in each lineage, but the transmembrane regions converged on a similar compositional solution centered around ATS. Our ATS engineering experiments supported this interpretation. Strong MTSs added to codon-optimized WT-hATP6 did not enable mitochondrial localization. However, even conservative ATS changes, chosen from natural ATP6 alignments and checked by structural modeling, were sufficient to improve mitochondrial localization of a highly hydrophobic protein. This emphasizes that both a capable MTS and appropriate hydrophobicity are required for mitochondrial targeting. These results build on earlier allotopic expression studies that focused on optimized MTS and codon usage, by indicating that TMH hydrophobicity is also an important determinant of mitochondrial targeting (Artika, 2020; Boominathan et al., 2016; Chin et al., 2018).

These pressures are not confined to nuclear-encoded proteins. The presence of ATS in both nuclear- and mtDNA-encoded IMM proteins suggests that similar pressures act on both groups of proteins. The high cost of aggregation-prone intermediates for mtDNA-encoded OXPHOS subunits could explain why these proteins also benefit from reduced TMH hydrophobicity, even without the challenge of cytosolic transit.

The challenge of this tradeoff in any species is likely a function of the insertion machinery. Organisms that retain alternative insertases would be expected to experience weaker solubility-stability constraints and consequently less demand for ATS. Plant and jakobid mitochondria retain Sec- or bacterial-type insertion components along with the Oxa1 pathway (Ghifari et al., 2018; Petrů et al., 2021), whereas most eukaryotes, including the metazoans and green algae examined here, rely on Oxa1 as the principal IMM insertase (Homberg et al., 2023). A Sec-type translocon is predicted to provide a protected conduit for the membrane insertion of more hydrophobic TMHs (Mercier et al., 2022), thereby buffering the cost of high hydrophobicity at the level of the machinery rather than the sequence. Where such additional pathways persist, the burden of solving both aqueous compatibility and membrane stability on the TMH sequence itself is likely reduced. Such comparisons argue that ATS is favored when targeting and insertion mechanisms make the solubility–stability tradeoff especially challenging.

These findings carry implications for both protein engineering and mitochondrial evolution. Many primary mitochondrial diseases arise from mutations affecting IMM proteins, for which gene therapy is the most likely curative strategy (Corrà et al., 2026). However, several attempts at gene therapy for mitochondrial disease have failed, likely due to the extreme hydrophobicity of mtDNA-encoded proteins (Artika, 2020). ATS may therefore offer a route toward functional nuclear genome expression as a therapeutic strategy for this devastating group of diseases. Evolutionarily, ATS provides a candidate sequence-level mechanism for addressing the aqueous compatibility barrier that has long been proposed to constrain transfer of mitochondrial genes to the nucleus. This framework may help explain why some hydrophobic mitochondrial genes have been successfully relocated during evolution, whereas highly hydrophobic OXPHOS subunits remain mitochondrially encoded in most lineages (Butenko et al., 2024). Our observations also point to a comparatively unexplored aspect of mitochondrial protein biogenesis, a field that has mostly focused on import machinery and targeting sequences while treating the sequence as relatively invariant. Beyond ATS itself, the features that govern mitochondrial membrane protein biology may be most readily discovered by learning from the solutions evolution has already found.

## Acknowledgments

We thank members of the Rutter lab for feedback and comments on the manuscript. We thank Prof. Sihem Boudina, Dr. Katsuhiko Funai, Dr. Peter S Shen, and Prof. Nels Elde for their guidance, and Dr. Nathan Krah, Dr. Alex Bott, Prof. Thomas Langer and Prof. Michal Minczuk for helpful discussions. Several figures were created with BioRender, or adapted from BioDraws, and from https://commons.wikimedia.org/wiki/File:Gauche-eclipsed_interconversion.svg under https://creativecommons.org/licenses/by/4.0/.

## Funding

This work was supported by the following funding:

American Heart Association 25PRE1372133 (TY)

National Science Foundation GRFP 2445150 (JB)

National Institutes of Health F32GM140525 (CNC)

National Institutes of Health National Institute of Diabetes and Digestive and Kidney Diseases 5T32DK091317 (JC)

American Heart Association 26PRE1557003 (JC)

National Institutes of Health R01GM137109 (MG)

National Institutes of Health R35GM131854 (JR)

JR is an investigator of the Howard Hughes Medical Institute.

## Author contributions

Conceptualization: TY, JR

Formal analysis: TY, JB, AJNP, BS, CNC, JC

Funding acquisition: JR

Investigation: TY, JB

Methodology: TY

Project administration: JR

Resources: JR, MG

Supervision: JR

Validation: TY

Visualization: TY, JB

Writing — original draft: TY, JB, MG, JR

Writing — review and editing: All authors

---

**Figure 1—figure supplement 1:**
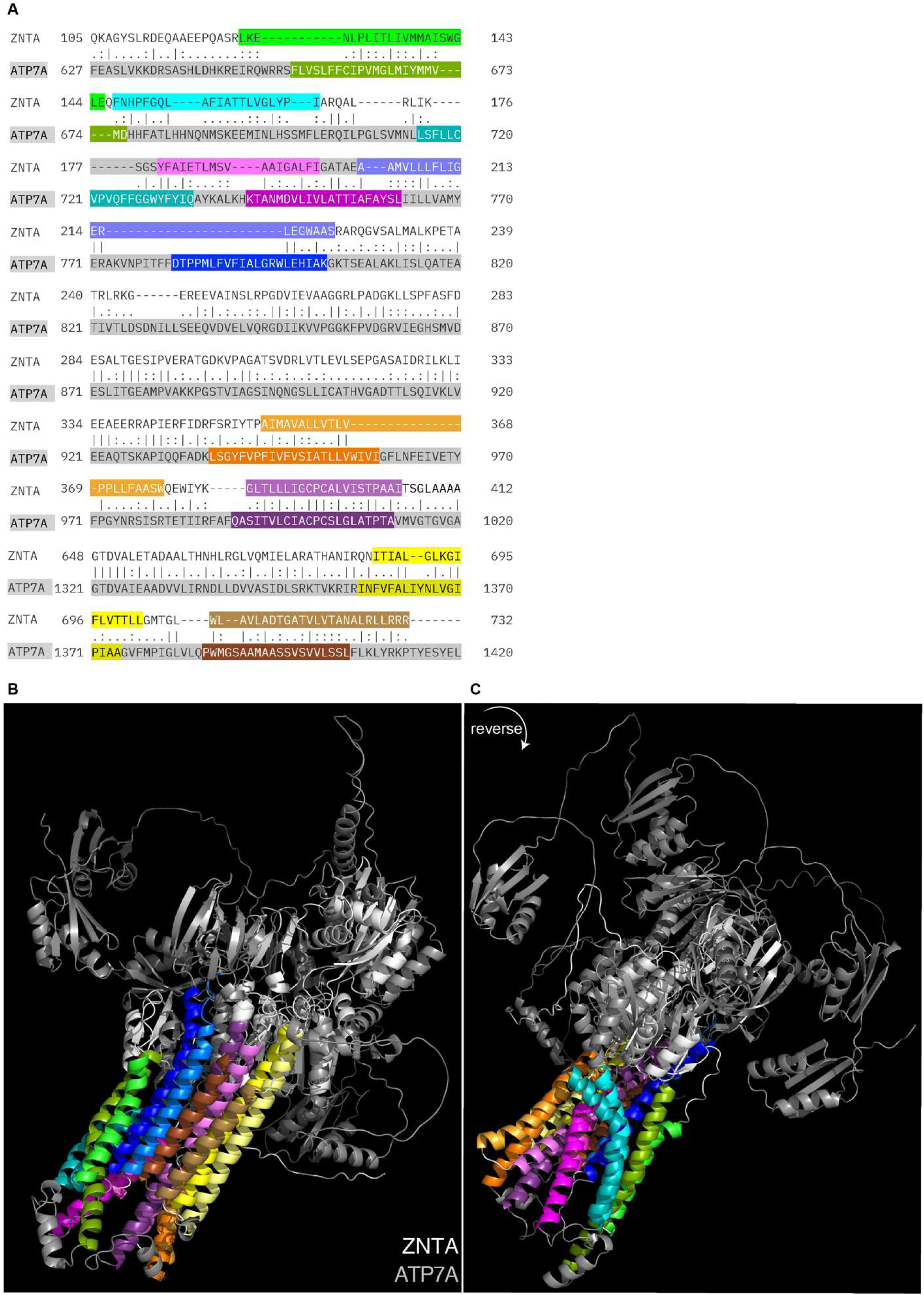
Structural alignment enables comparison of divergent proteins. **(A)** Needleman-Wunsch sequence alignment of *E. coli* ZNTA and human ATP7A. **(B, C)** PyMOL structural alignment (RMSD = 2.373 Å) shown in two orientations. Transmembrane helices are colored consistently across all panels; lighter and darker shades denote *E. coli* and human sequences respectively.

**Figure 2—figure supplement 1:**
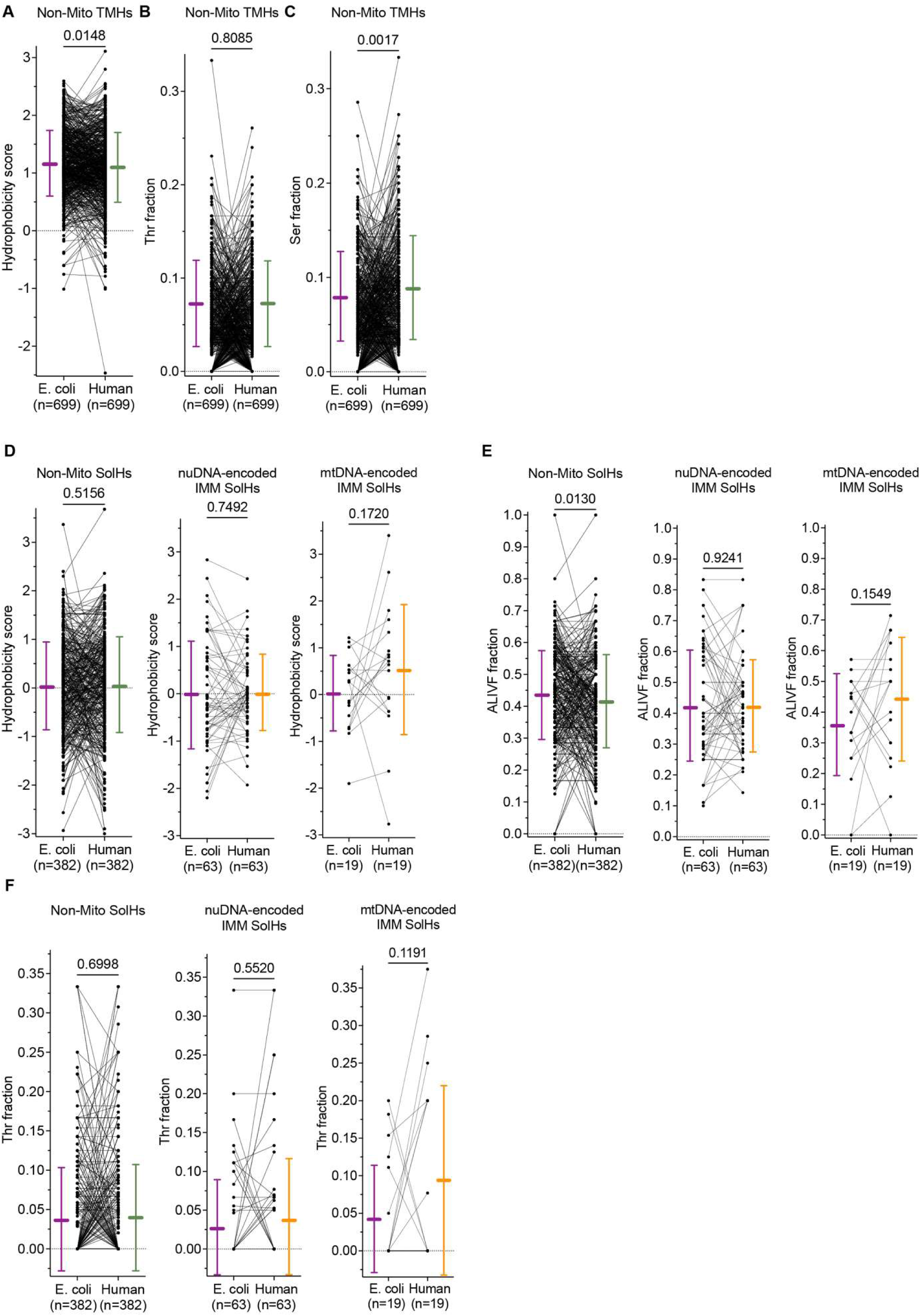
ATS is specific to mitochondrial TMHs over evolution. **(A–C)** Hydrophobicity score (A), threonine fraction (B), and serine fraction (C) of structurally aligned TMH pairs from evolutionarily related *E. coli* and *H. sapiens* non-IMM TM proteins. **(D–F)** Hydrophobicity score (D), aliphatic residue fraction (E), and threonine fraction (F) of structurally aligned SolH pairs from evolutionarily related *E. coli* and *H. sapiens* non-mitochondrial, nuDNA-encoded IMM, and mtDNA-encoded IMM TM proteins. Lines connect paired helices; bars represent mean ± SD. p-values from Wilcoxon signed-rank test.

**Figure 2—figure supplement 2:**
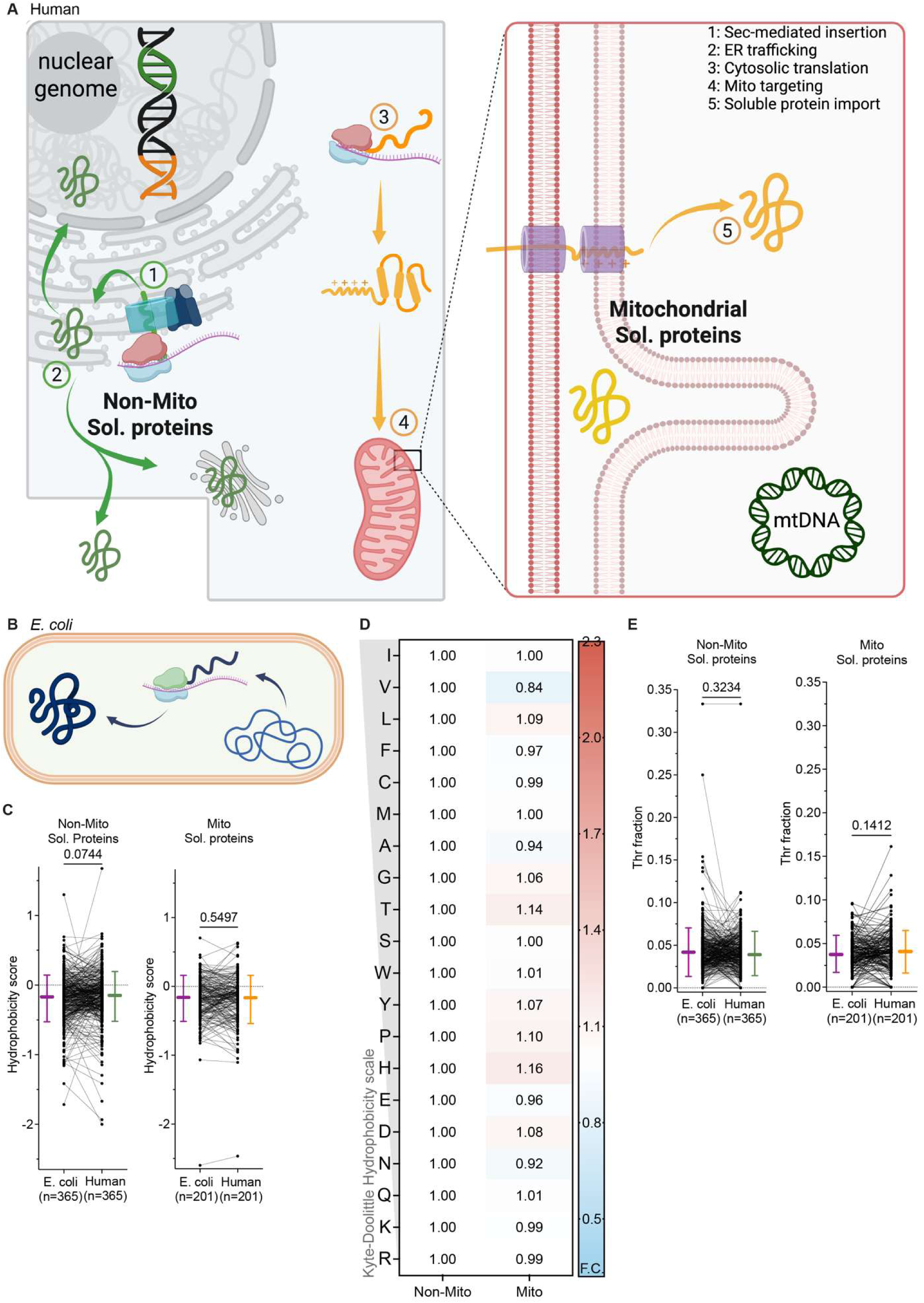
Mitochondrial soluble proteins do not show ATS relative to proteobacterial counterparts. **(A)** Non-mitochondrial soluble proteins are synthesized and targeted to various compartments (1, 2), bypassing solubility constraints. Nuclear-encoded mitochondrial soluble proteins are post-translationally imported into the matrix (3, 4) and released as soluble proteins (5). Unlike inner mitochondrial membrane TM proteins, neither class of soluble proteins faces hydrophobicity-driven solubility constraints. **(B)** *E. coli* soluble proteins are translated and folded in the cytoplasm, bypassing hydrophobicity-driven solubility constraints. **(C)** Hydrophobicity score of structurally aligned soluble helix pairs from evolutionarily related *E. coli* and *H. sapiens* non-mitochondrial and mitochondrial soluble proteins. **(D)** Mean amino acid composition of helices from evolutionarily related *H. sapiens* mitochondrial soluble proteins, expressed as fold change relative to paired *E. coli* helices and normalized to the corresponding fold change in non-mitochondrial soluble proteins. Red indicates enrichment, blue indicates depletion. **(E)** Related to (C), threonine fraction of structurally aligned soluble helix pairs from evolutionarily related *E. coli* and *H. sapiens* non-mitochondrial and mitochondrial soluble proteins. Lines connect paired helices; bars represent mean ± SD. p-values from Wilcoxon signed-rank test.

**Figure 2—figure supplement 3:**
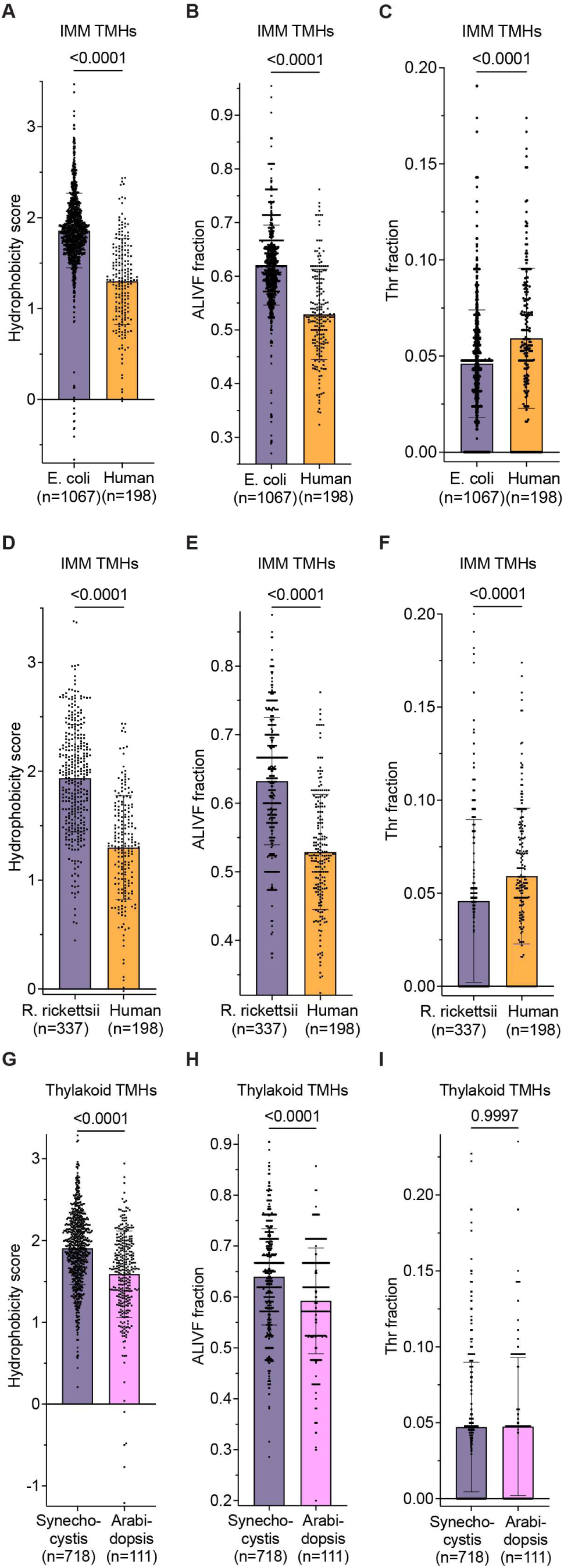
Mitochondrial, but not chloroplast thylakoid, membrane proteins are threonine-enriched relative to bacterial counterparts. **(A–C)** Hydrophobicity score (A), ALIVF fraction (B), and threonine fraction (C) of all UniProtKB-annotated TMHs from *E. coli* and *H. sapiens* IMM-targeted TM proteins. **(D–F)** Hydrophobicity score (D), ALIVF fraction (E), and threonine fraction (F) of all UniProtKB-annotated TMHs from *R. rickettsii* and *H. sapiens* IMM-targeted TM proteins. **(G–I)** Hydrophobicity score (G), ALIVF fraction (H), and threonine fraction (I) of all UniProtKB-annotated TMHs from *Synechocystis* sp. and *A. thaliana* chloroplast thylakoid membrane proteins. Bars represent mean ± SD. p-values from a Mann–Whitney U test.

**Figure 2—figure supplement 4:**
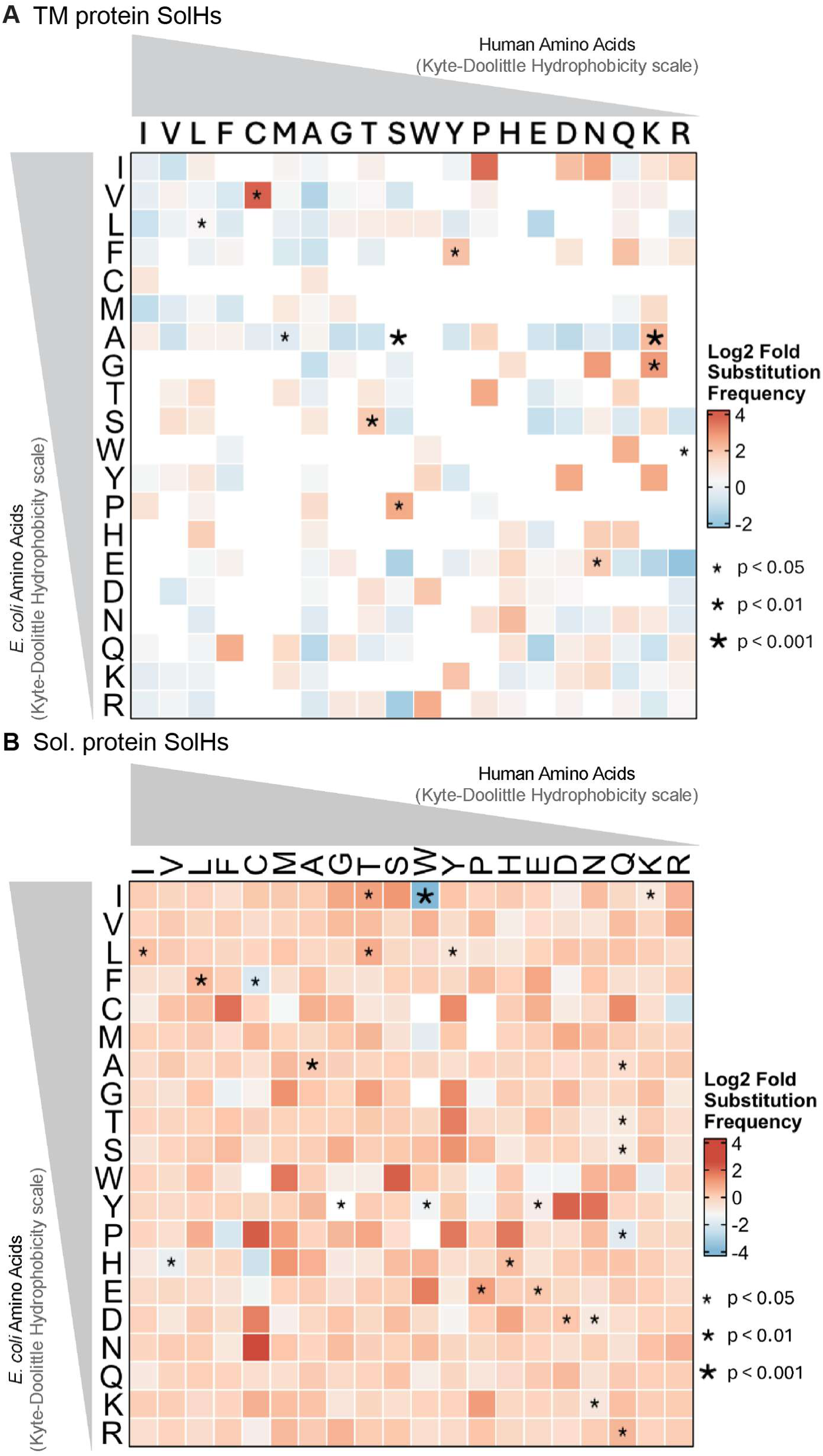
Substitution matrices for soluble helix controls show no ATS enrichment. **(A)** Relative substitution matrix for aligned SolHs from evolutionarily related *E. coli* and *H. sapiens* IMM TM proteins, normalized to substitutions from non-IMM SolHs. **(B)** Relative substitution matrix for aligned helices from evolutionarily related *E. coli* and *H. sapiens* soluble proteins, normalized to substitutions from non-mitochondrial soluble proteins. Asterisks denote statistical significance from Fisher’s exact test (*p<0.05, **p<0.01, ***p<0.001).

**Figure 3—figure supplement 1:**
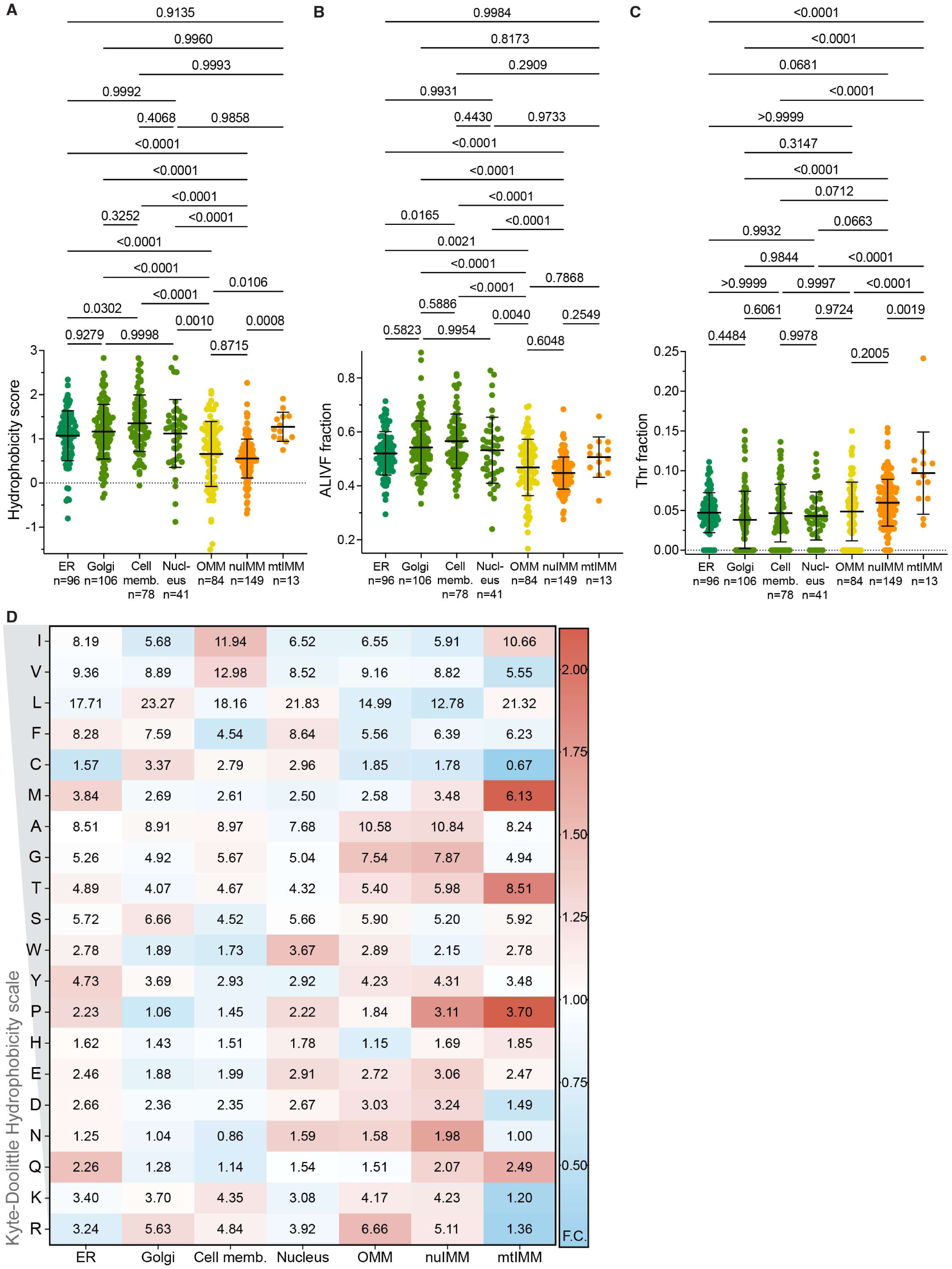
Structurally annotated mitochondrial TMHs have lower hydrophobicity and higher threonine content. **(A–C)** Hydrophobicity scores (A), ALIVF fraction (B), and threonine fraction (C) of human TMHs identified from TM protein AlphaFold2 structures, grouped by membrane compartment, with all pairwise comparisons shown. Each point represents one TM protein; bars represent mean ± SD. p-values from one-way ANOVA with Tukey’s post-hoc test. **(D)** Amino acid residue frequency (%) in human TMHs identified from TM protein AlphaFold structures, grouped by membrane compartment. Cell values represent mean residue frequency; color scale reflects fold-change relative to the mean of non-mitochondrial compartments (red, enriched; blue, depleted).

**Figure 3—figure supplement 2:**
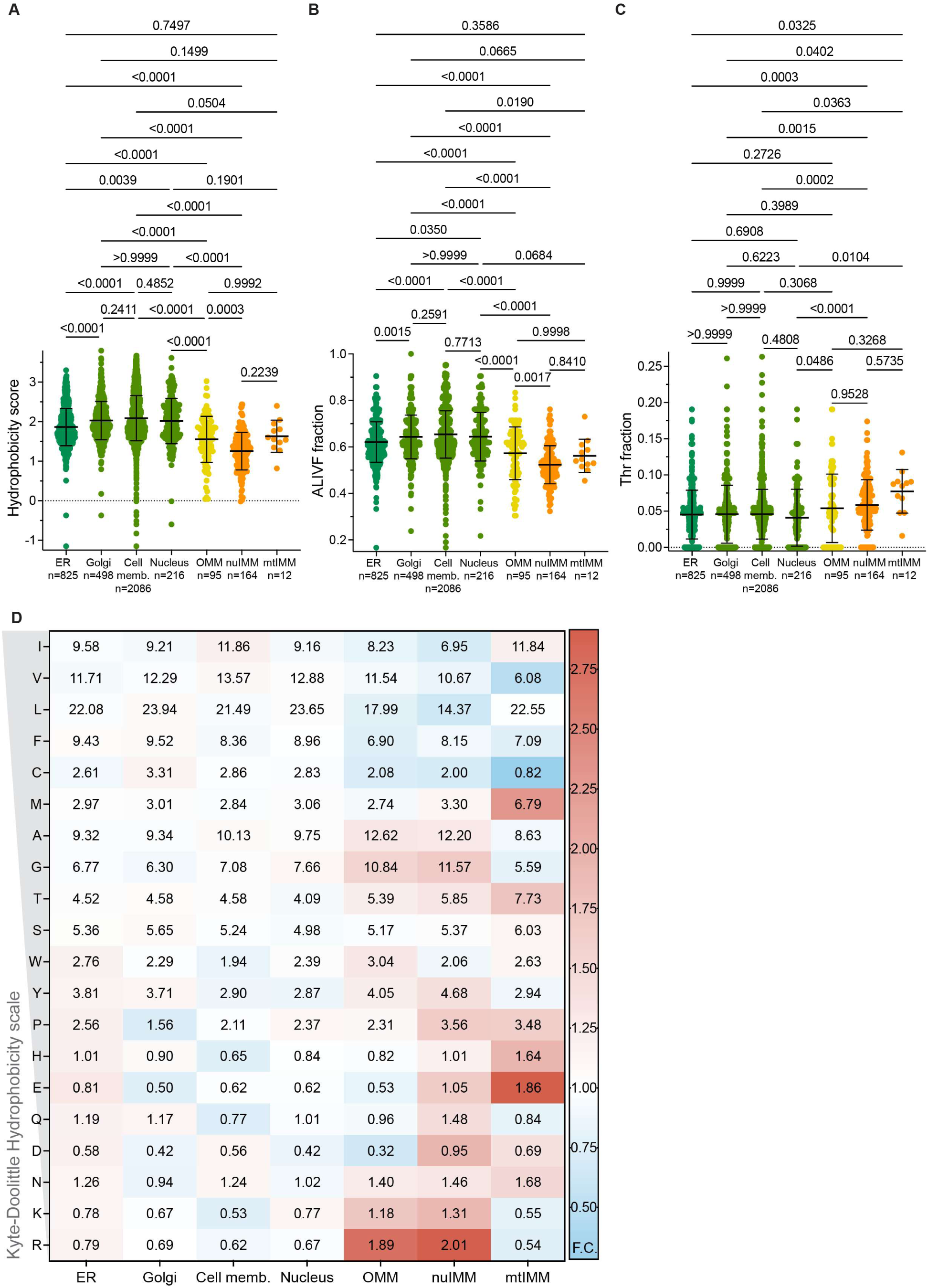
UniProtKB-annotated mitochondrial transmembrane helices have lower hydrophobicity and higher threonine. **(A–C)** Hydrophobicity scores (A), ALIVF fraction (B), and threonine fraction (C) of human TMHs identified from TM protein UniProtKB annotations, grouped by membrane compartment. Each point represents one TM protein; bars represent mean ± SD. p-values from one-way ANOVA with Tukey’s post-hoc test. **(D)** Amino acid residue frequency (%) in human TMHs identified from TM protein UniProtKB annotations, grouped by membrane compartment. Cell values represent mean residue frequency; color scale reflects fold-change relative to the mean of non-mitochondrial compartments (red, enriched; blue, depleted).

**Figure 3—figure supplement 3:**
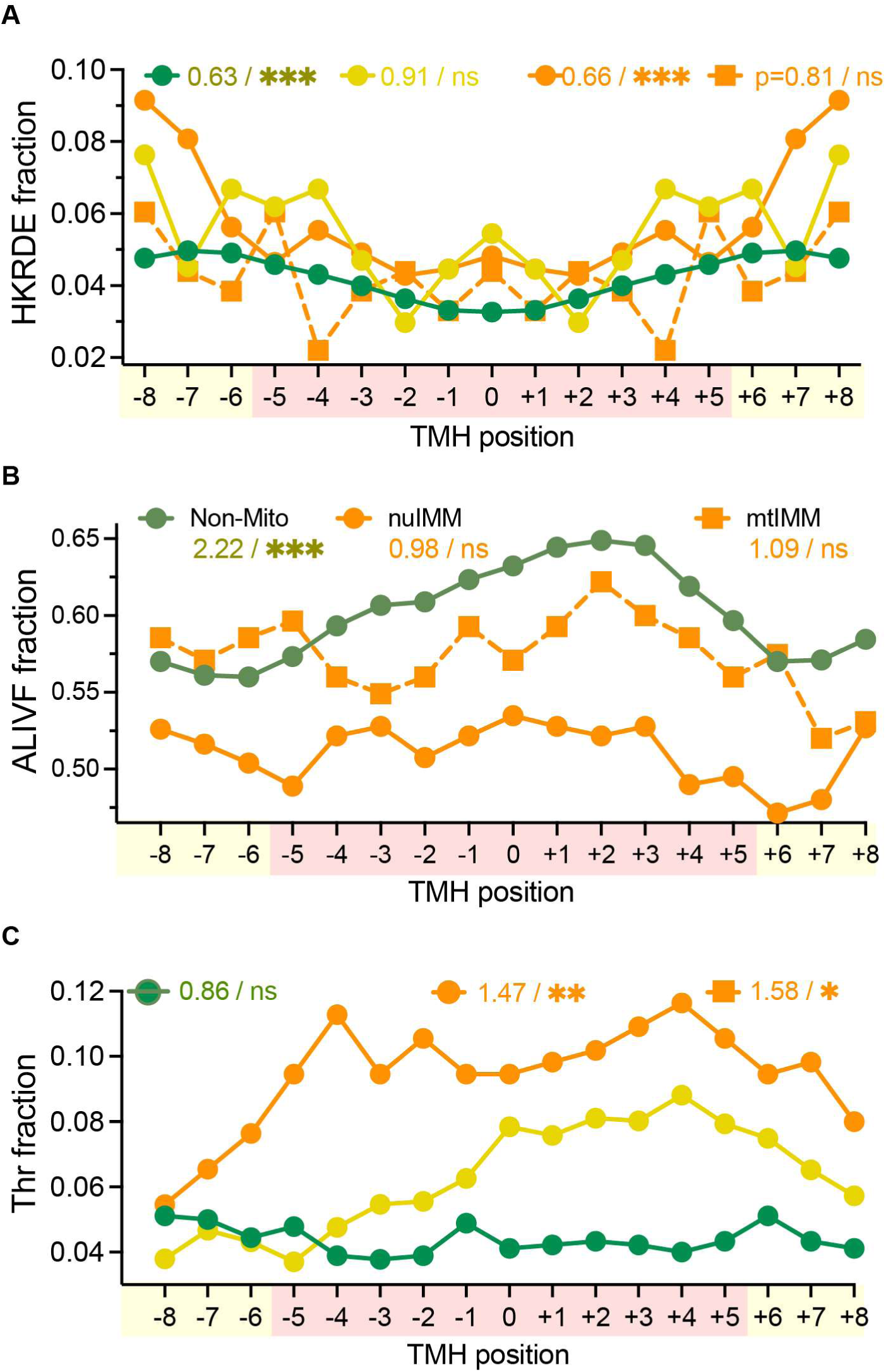
Threonine is enriched in the core of mitochondrial TMHs. **(A)** Positional distribution of HKRDE fraction along TMHs from human TM proteins, aligned agnostically to membrane orientation, grouped by membrane compartment. **(B, C)** Positional distribution of ALIVF fraction (B) and threonine fraction (C) along a separate set of TMHs oriented by structurally informed annotation, grouped by membrane compartment. Values on figure denote odds ratio/significance for core vs. peripheral enrichment from Fisher’s exact test (✱p<0.05, ✱✱p<0.01, ✱✱✱p<0.001).

**Figure 3—figure supplement 4:**
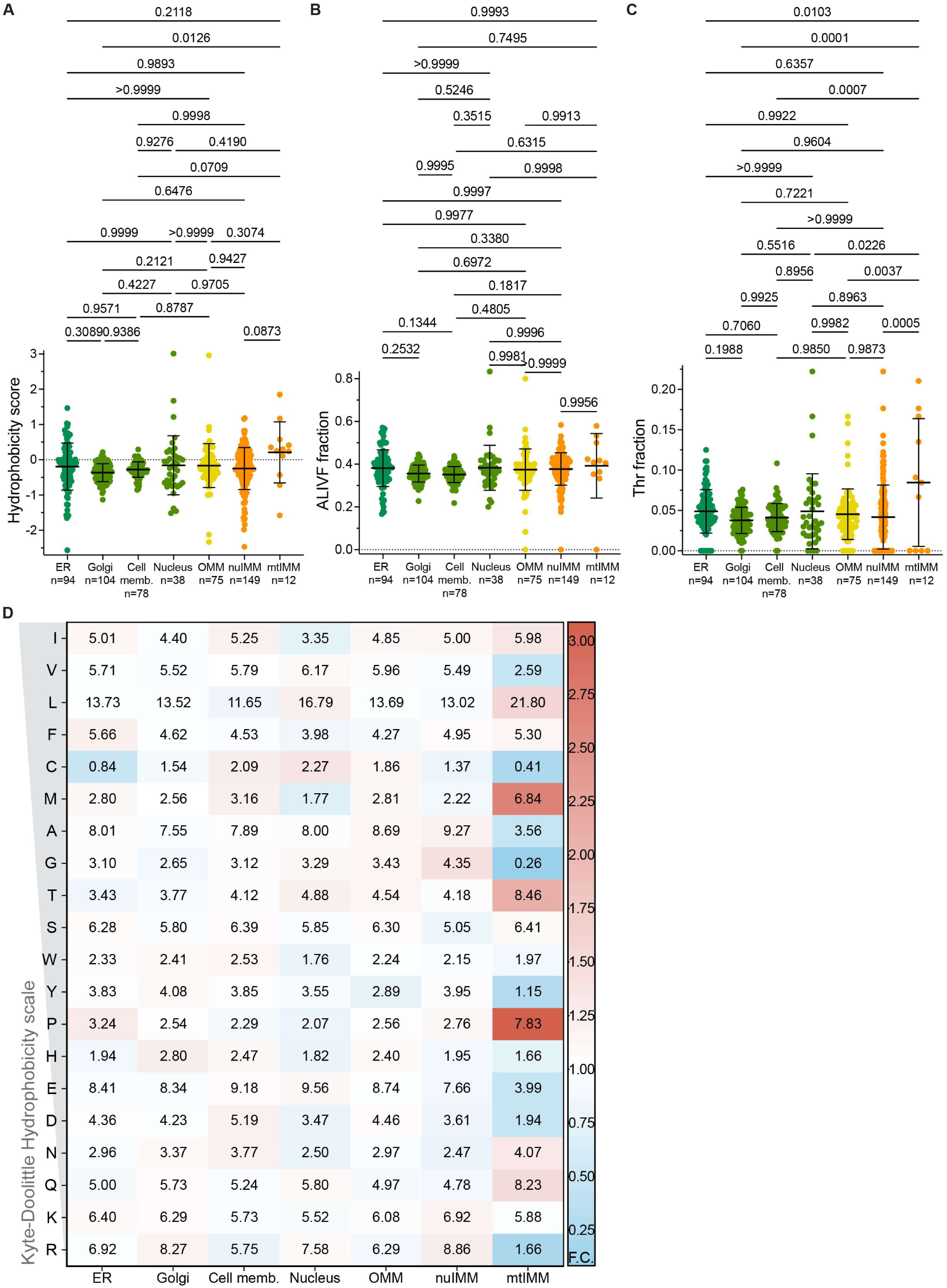
Soluble helices of nuclear-encoded TM proteins have similar hydrophobicity and composition. **(A–C)** Hydrophobicity scores (A), ALIVF fraction (B), and threonine fraction (C) of human soluble helices identified from TM protein AlphaFold2 structures, grouped by membrane compartment, with all pairwise comparisons shown. Each point represents one TM protein; bars represent mean ± SD. p-values from one-way ANOVA with Tukey’s post-hoc test. **(D)** Amino acid residue frequency (%) in human soluble helices identified from TM protein AlphaFold structures, grouped by membrane compartment. Cell values represent mean residue frequency; color scale reflects fold-change relative to the mean of non-mitochondrial compartments (red, enriched; blue, depleted).

**Figure 3—figure supplement 5:**
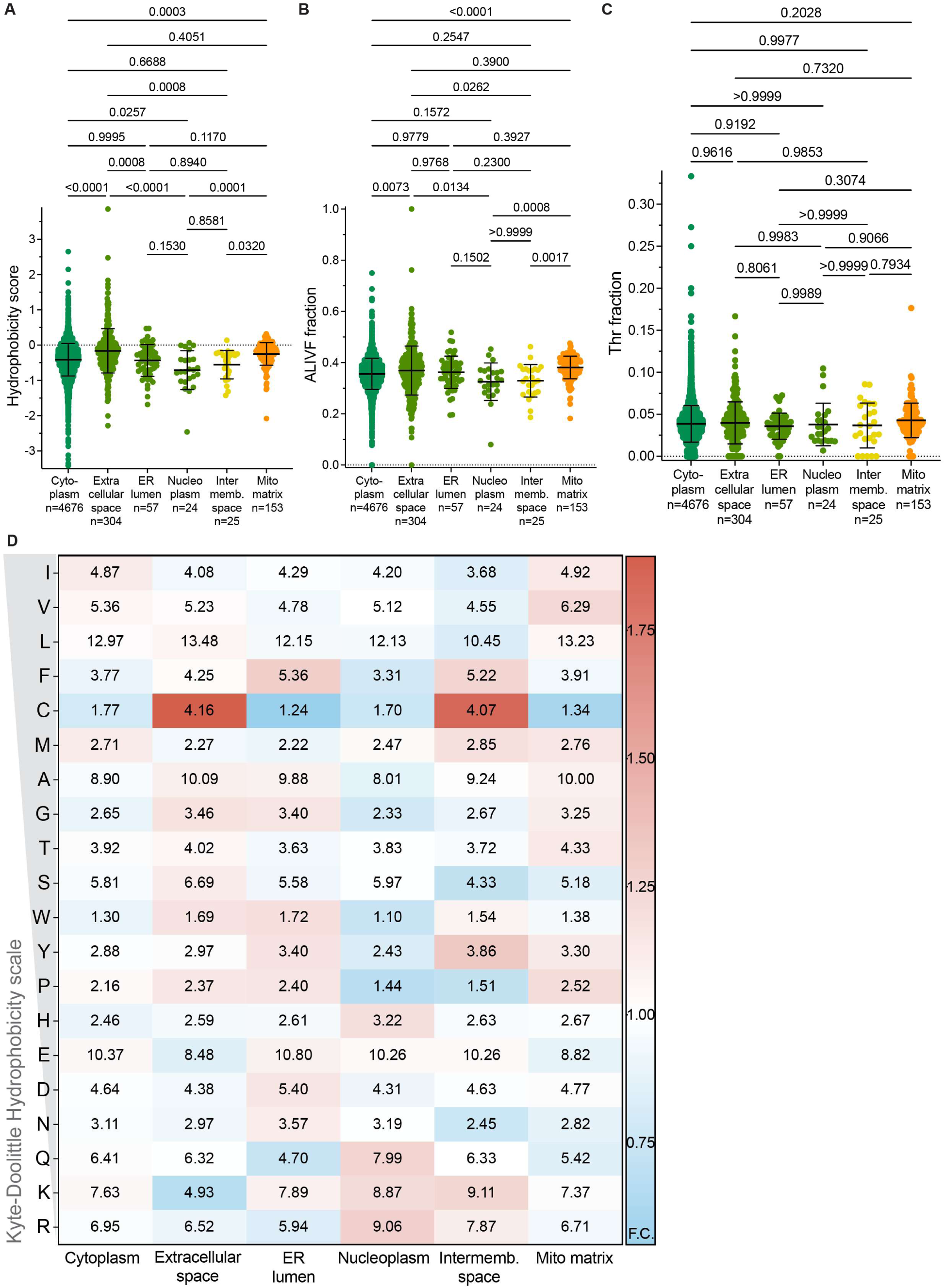
Soluble mitochondrial proteins do not show decreased hydrophobicity or altered composition. **(A–C)** Hydrophobicity scores (A), ALIVF fraction (B), and threonine fraction (C) of human soluble helices identified from soluble protein AlphaFold structures, grouped by compartment. Each point represents one soluble protein; bars represent mean ± SD. p-values from one-way ANOVA with Tukey’s post-hoc test. **(D)** Amino acid residue frequency (%) in human soluble helices identified from soluble protein AlphaFold2 structures, grouped by compartment. Cell values represent mean residue frequency; color scale reflects fold-change relative to the mean of non-mitochondrial compartments (red, enriched; blue, depleted).

**Figure 3—figure supplement 6:**
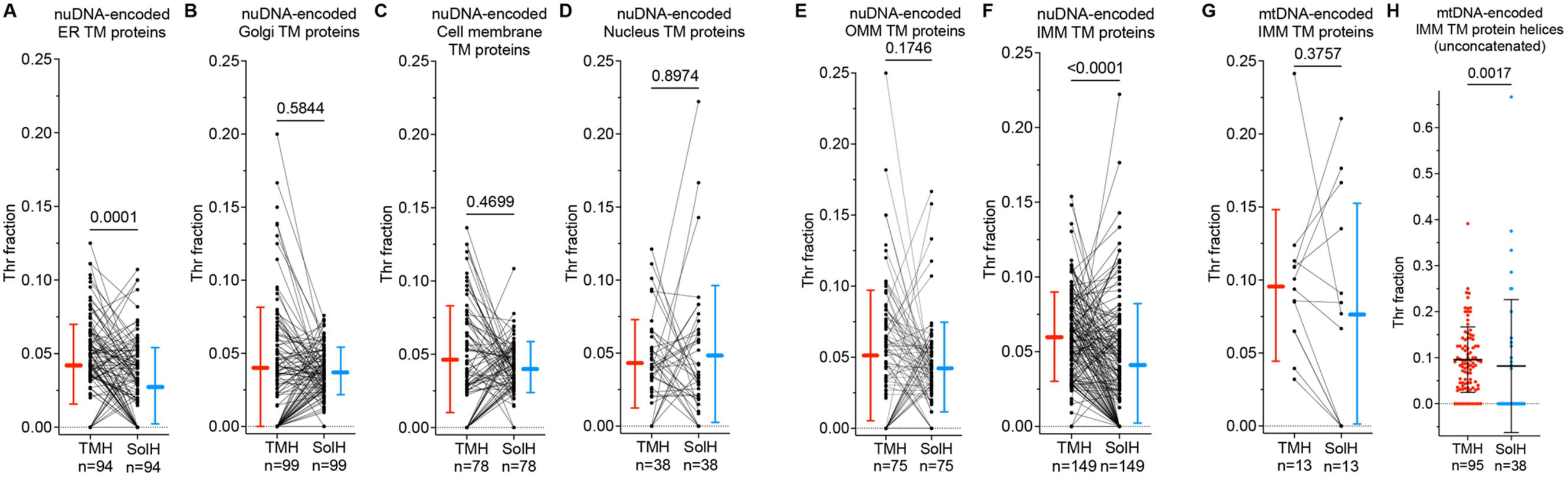
Threonine is enriched in transmembrane over soluble helices in IMM proteins. Threonine fraction of concatenated transmembrane versus soluble helices within each TM protein. **(A–D)** nuDNA-encoded non-mitochondrial TM proteins **(E, F)** nuDNA-encoded mitochondrial TM proteins **(G)** mtDNA-encoded IMM TM proteins **(H)** Threonine fraction of individual, unconcatenated helices from mtDNA-encoded IMM TM proteins Lines connect paired transmembrane and soluble helices from the same protein; bars represent mean ± SD. p-values from Wilcoxon signed-rank test (A–G) and Mann–Whitney test (H).

**Figure 3—figure supplement 7:**
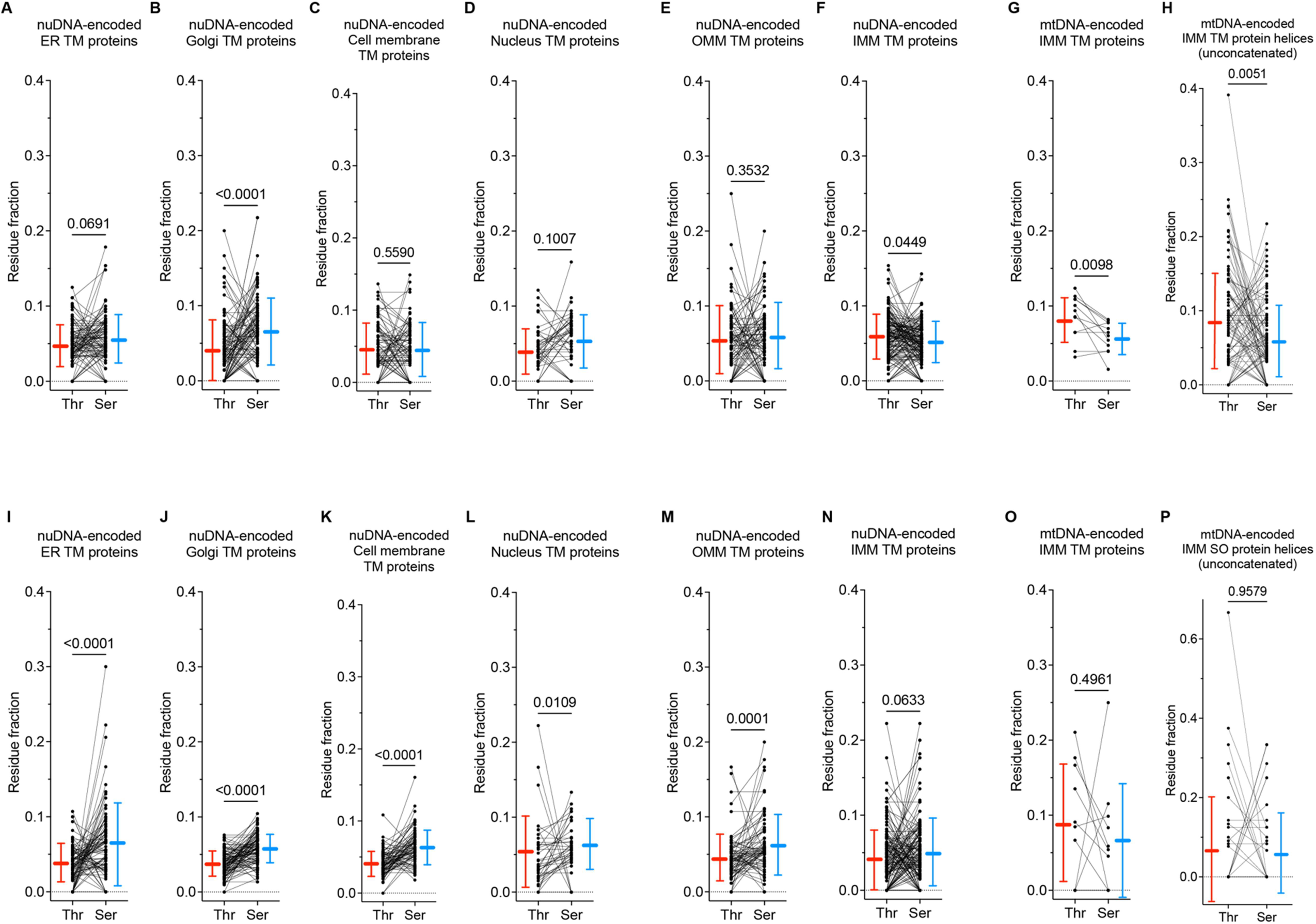
Serine preference over threonine is reversed in IMM transmembrane helices. Threonine versus serine residue fraction in helices from TM proteins. Top panels (A–H) show transmembrane helices; bottom panels (I–P) show soluble helices. Lines connect paired transmembrane and soluble helices from the same protein; bars represent mean ± SD. p-values from Wilcoxon signed-rank tests and Mann–Whitney tests for unconcatenated helices.

**Figure 3—figure supplement 8:**
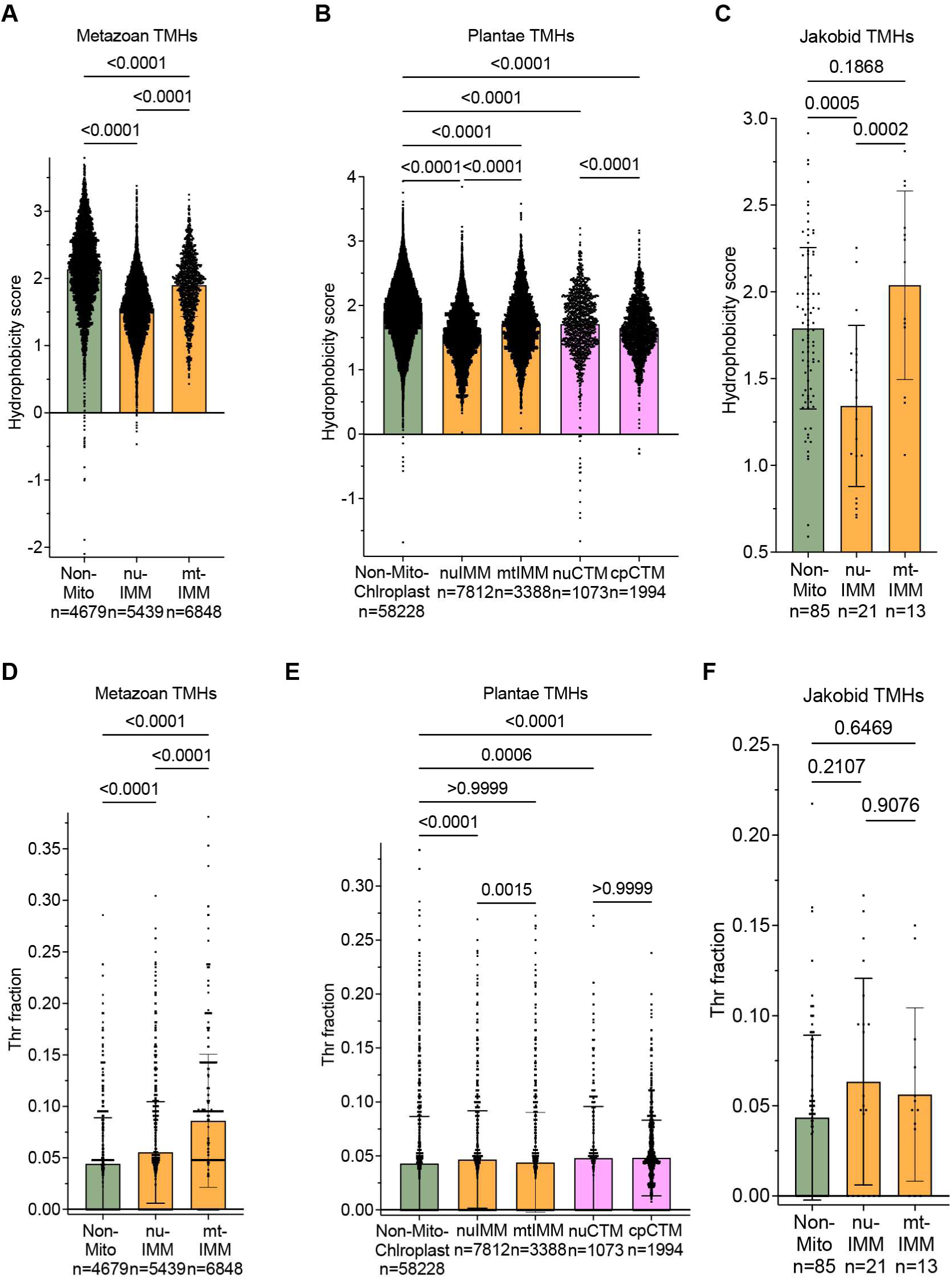
Targeting and insertion constraints shape eukaryotic TMH hydrophobicity and threonine content. **(A–C)** TMH hydrophobicity scores for metazoan (A), plant (B), and jakobid (C) proteins. **(D–F)** Threonine fractions within TMHs for the corresponding metazoan (D), plant (E), and jakobid (F) proteins. Proteins were grouped into non-mitochondrial, nuclear-encoded inner mitochondrial membrane, and mitochondrially encoded IMM classes; plant proteins additionally included nuclear-encoded and chloroplast genome-encoded chloroplast thylakoid membrane (CTM) proteins. Statistical significance was assessed using one-way ANOVA with multiple-comparison testing; adjusted p-values are shown above comparisons. Sample sizes are indicated below each group.

**Figure 4—figure supplement 1:**
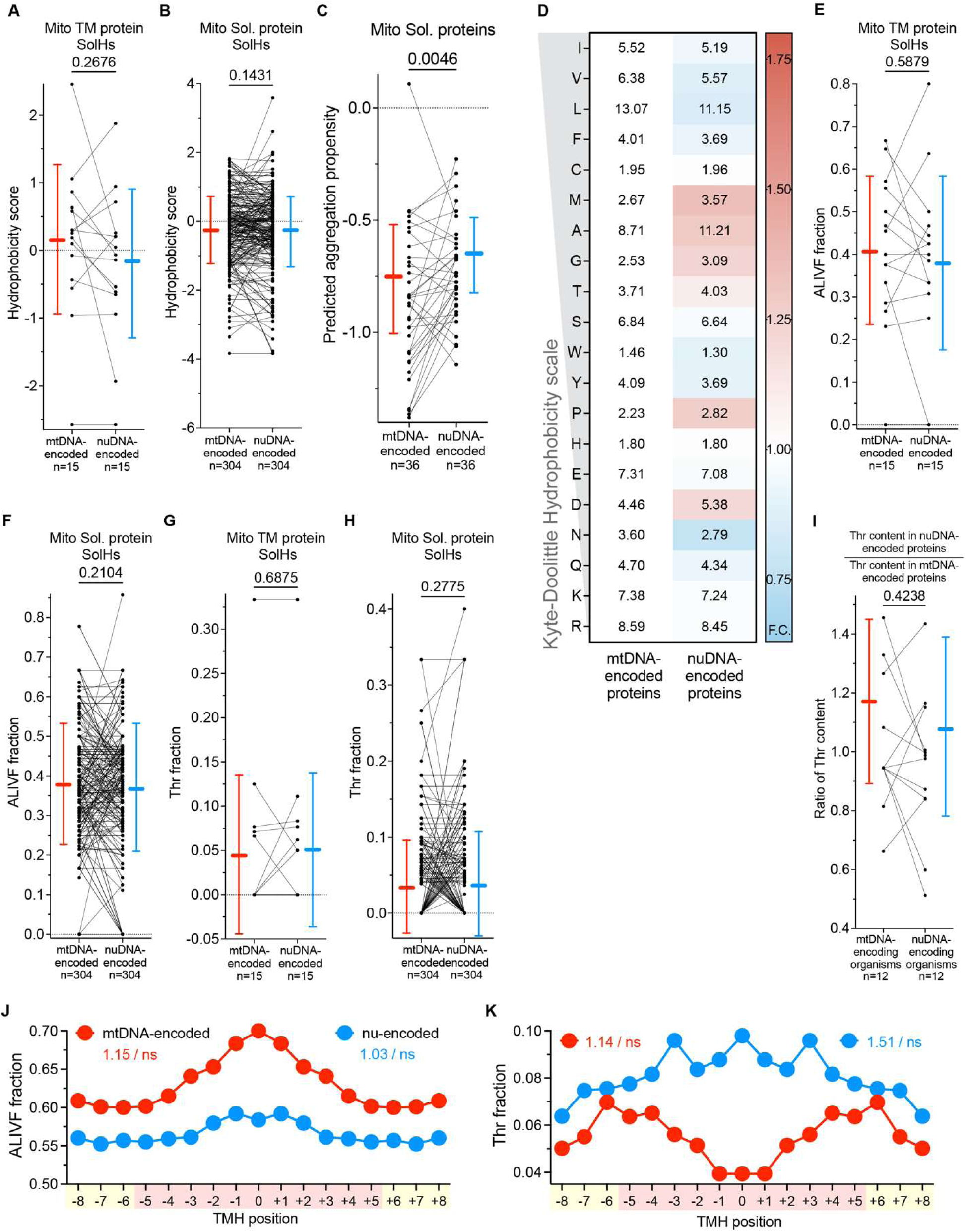
ATS is TMH-specific during mitochondrial gene transfers. **(A)** Hydrophobicity scores of structurally aligned soluble helix pairs from mtDNA- and nuDNA-encoded versions of the same IMM protein across eukaryotes. **(B)** Hydrophobicity scores of structurally aligned soluble helix pairs from mtDNA- and nuDNA-encoded versions of the same soluble mitochondrial protein across eukaryotes. **(C)** Predicted aggregation propensity of the above soluble proteins. **(D)** Amino acid residue frequency (%) in SolHs of mtDNA- and nuDNA-encoded soluble proteins across eukaryotes. Cell values represent mean residue frequency; color scale reflects fold-change relative to mtDNA-encoded proteins (red, enriched; blue, depleted). **(E–H)** ALIVF fraction **(E, F)** and threonine fraction **(G, H)** of structurally aligned soluble helix pairs from mtDNA- and nuDNA-encoded versions of the same IMM transmembrane protein **(E, G)** and soluble protein **(F, H)** across eukaryotes. Lines connect paired helices; bars represent mean ± SD. p-values from Wilcoxon signed-rank test. **(I)** Ratio of mean TMH threonine content in nuDNA- to mtDNA-encoded TM proteins, calculated separately for organisms retaining mitochondrial gene encoding and those having transferred the gene to the nuclear genome. Lines connect paired organisms; bars represent mean ± SD. p-value from Wilcoxon signed-rank test. **(J, K)** Positional distribution of ALIVF fraction (J) and threonine fraction (K) along UniProtKB-annotated TMHs from mtDNA- and nuDNA-encoded IMM proteins across eukaryotes aligned agnostically to membrane orientation. Values on figure denote odds ratio/significance for core vs. peripheral enrichment from Fisher’s exact test (✱p<0.05, ✱✱p<0.01, ✱✱✱p<0.001).

**Figure 5—figure supplement 1:**
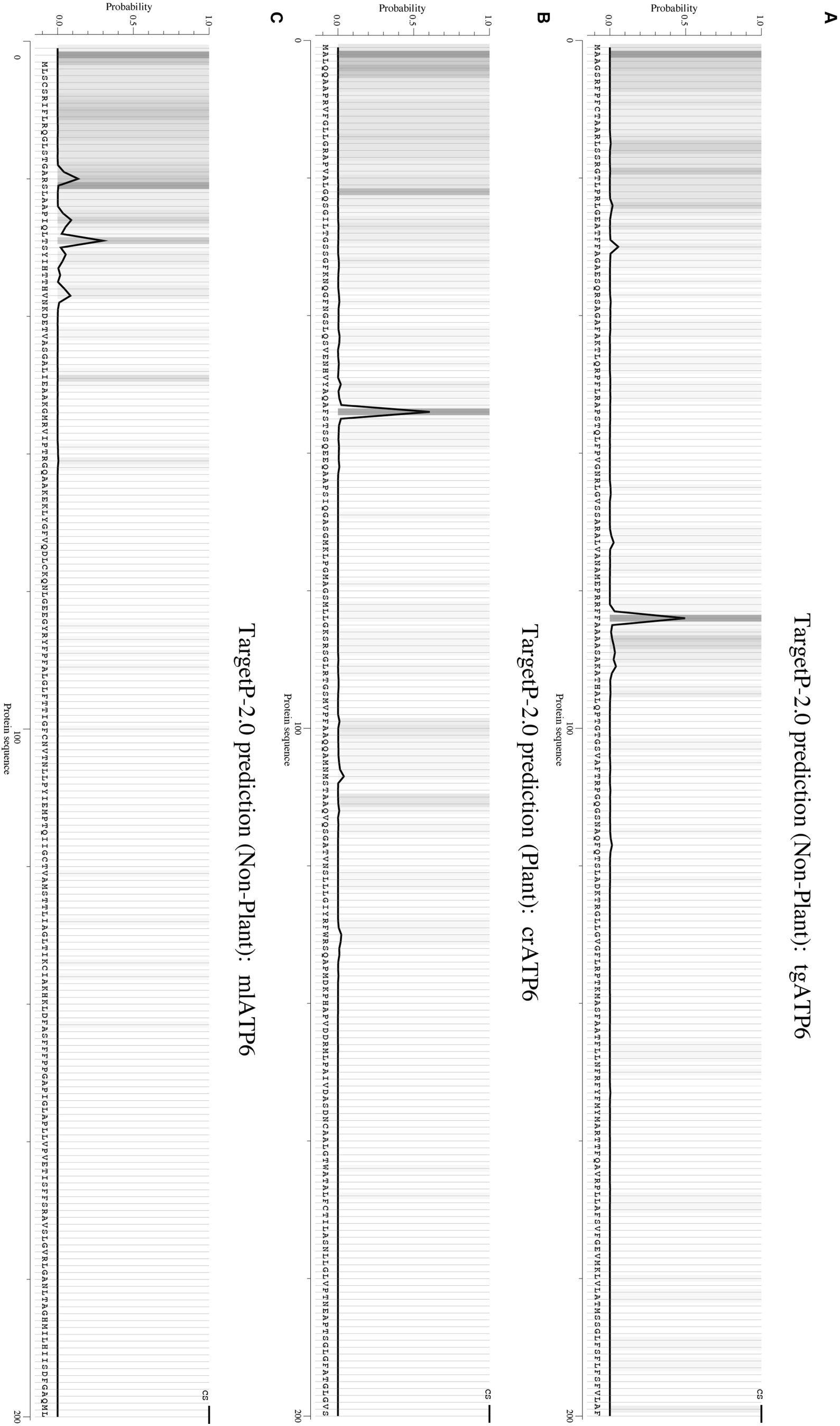
Nuclear-encoded ATP6s possess cleavable N-terminal MTSs. **(A–C)** TargetP 2.0 prediction plots for nuclear-encoded ATP6 sequences. The y-axis shows predicted probability of mitochondrial targeting sequence (MTS) identity at each residue position. The predicted cleavage site (CS) is indicated. (A) mlATP6, (B) crATP6, (C) tgATP6. Predicted MTS likelihoods: mlATP6 = 0.99, crATP6 = 0.56, tgATP6 = 0.49.

**Figure 5—figure supplement 2:**
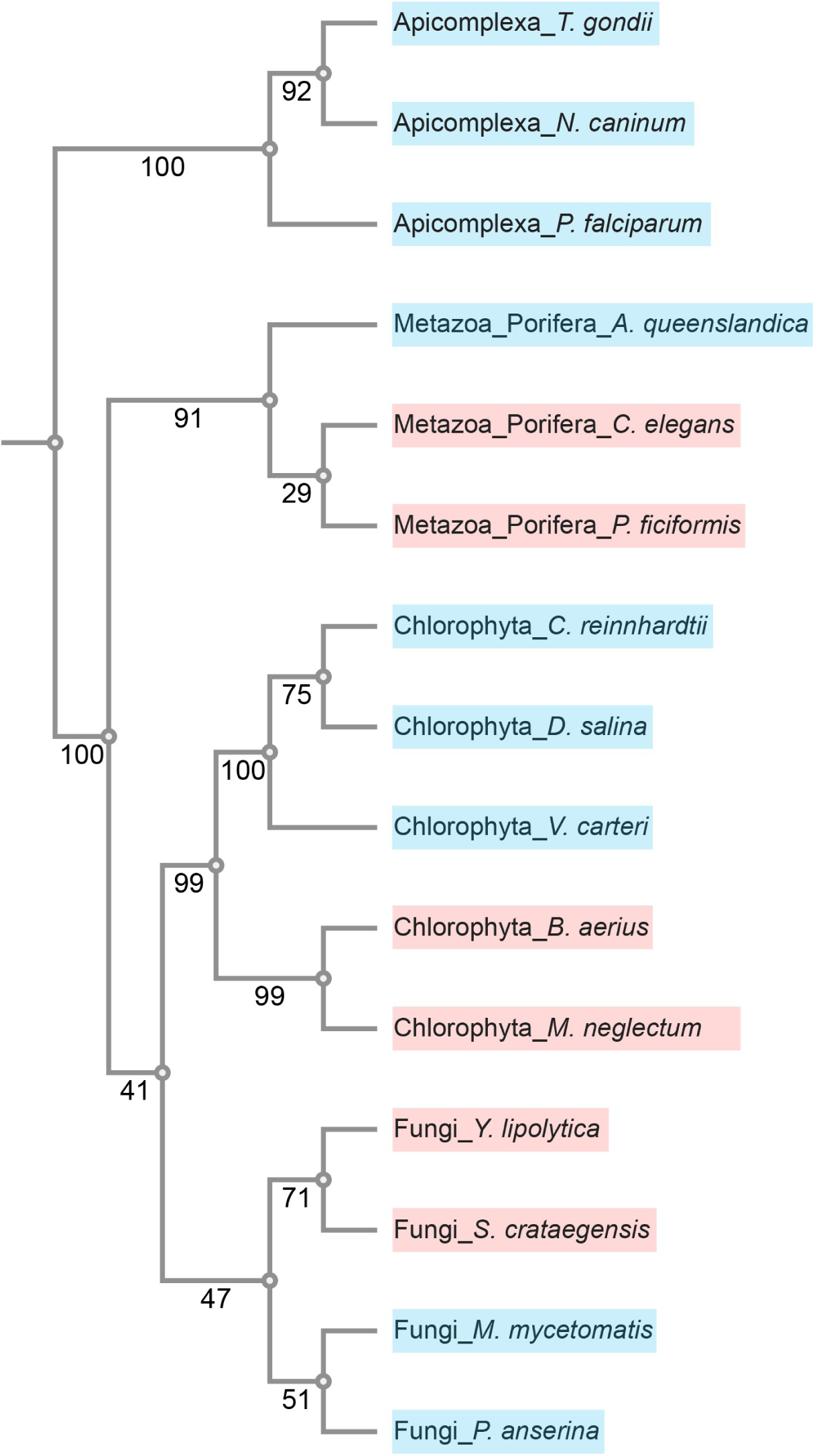
*MT-ATP9* underwent independent nuclear gene transfers. Phylogenetic tree of MT-ATP9 sequences. Blue indicates nuclear-encoded ATP9; red indicates mitochondrially encoded ATP9. Numbers at nodes indicate bootstrap support values.

**Figure 5—figure supplement 3:**
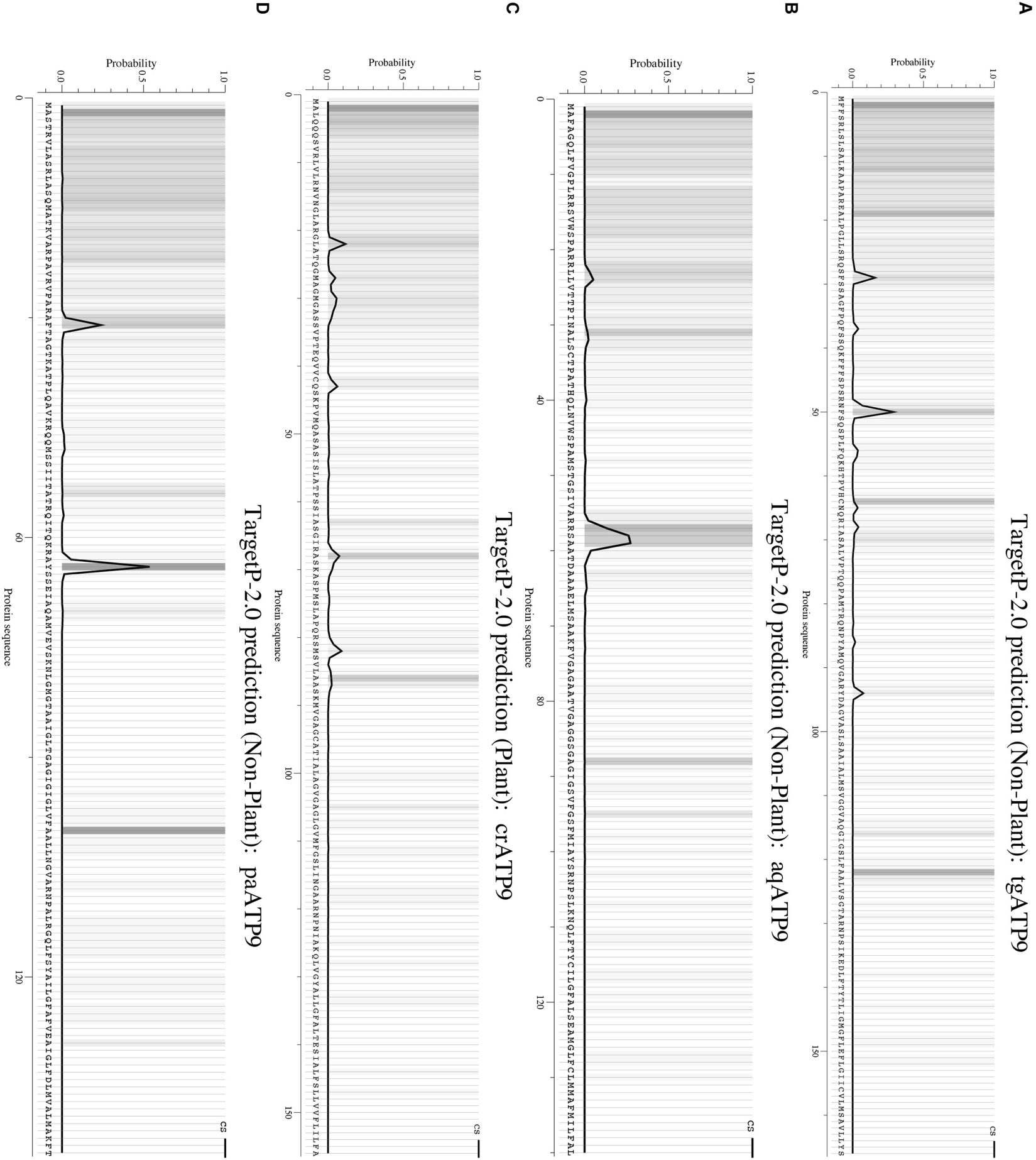
Nuclear-encoded ATP9s possess cleavable N-terminal MTSs. **(A–D)** TargetP 2.0 prediction plots for nuclear-encoded ATP9 sequences. The y-axis shows predicted probability of mitochondrial targeting sequence (MTS) identity at each residue position. The predicted cleavage site (CS) is indicated. (A) tgATP9 (B) aqATP9 (C) crATP9 (D) paATP9; Predicted MTS likelihoods: tgATP9 = 0.94, aqATP9 = 0.96, crATP9 = 0.59, paATP9 = 0.99.

**Figure 5—figure supplement 4:**
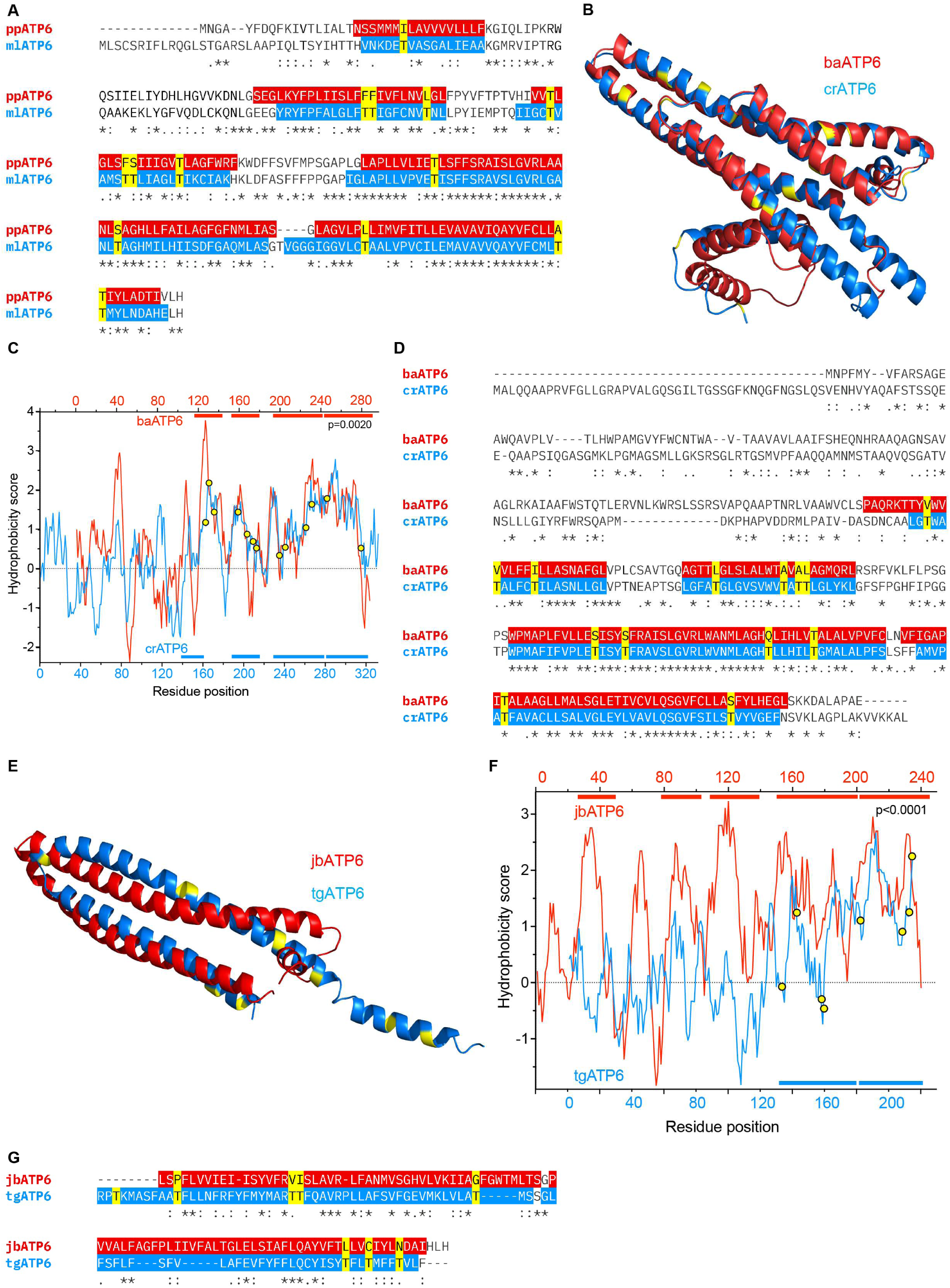
Sequence and structural alignment of ATP6s shows ATS. **(A)** ClustalW sequence alignment of ppATP6 and mlATP6. Red and blue highlights indicate UniProtKB-annotated transmembrane helices in ppATP6 and mlATP6, respectively. Yellow indicates threonine residues in mlATP6 TMHs. **(B)** Structural alignment of crATP6 (blue) and baATP6 (red) reveals conservation of proton channel architecture following nuclear gene transfer. Yellow indicates threonine residues in crATP6. RMSD = 0.868 Å. The putative MTS and TMH1 were omitted from the structural visualization as they do not form part of the proton channel architecture. **(C)** Whole-protein hydrophobicity profiles of crATP6 (blue) and baATP6 (red). Colored bars above and below indicate predicted transmembrane helices for baATP6 and crATP6, respectively. Yellow circles indicate threonine residues in crATP6 TMHs. p<0.0001, Wilcoxon rank-sum test comparing TMH hydrophobicity scores between crATP6 and baATP6. **(D)** ClustalW sequence alignment of baATP6 and crATP6. Red and blue highlights indicate UniProtKB-annotated transmembrane helices in baATP6 and crATP6, respectively. Yellow indicates threonine residues in crATP6 TMHs. **(E)** Structural alignment of tgATP6 (blue) and jbATP6 (red) reveals conservation of proton channel architecture following nuclear gene transfer. Yellow indicates threonine residues in tgATP6. RMSD = 10.109 Å. TMHs 1–3 of jbATP6 were omitted from the structural visualization as they do not form part of the proton channel architecture. **(F)** Whole-protein hydrophobicity profiles of tgATP6 (blue) and jbATP6 (red). Colored bars above and below indicate predicted transmembrane helices for jbATP6 and tgATP6, respectively. Yellow circles indicate threonine residues in tgATP6 TMHs. p<0.0001, Wilcoxon rank-sum test comparing TMH hydrophobicity scores between tgATP6 and jbATP6. **(G)** ClustalW sequence alignment of jbATP6 and tgATP6. Red and blue highlights indicate UniProtKB-annotated transmembrane helices in jbATP6 and tgATP6, respectively. Yellow indicates threonine residues in tgATP6 TMHs.

**Figure 6—figure supplement 1:**
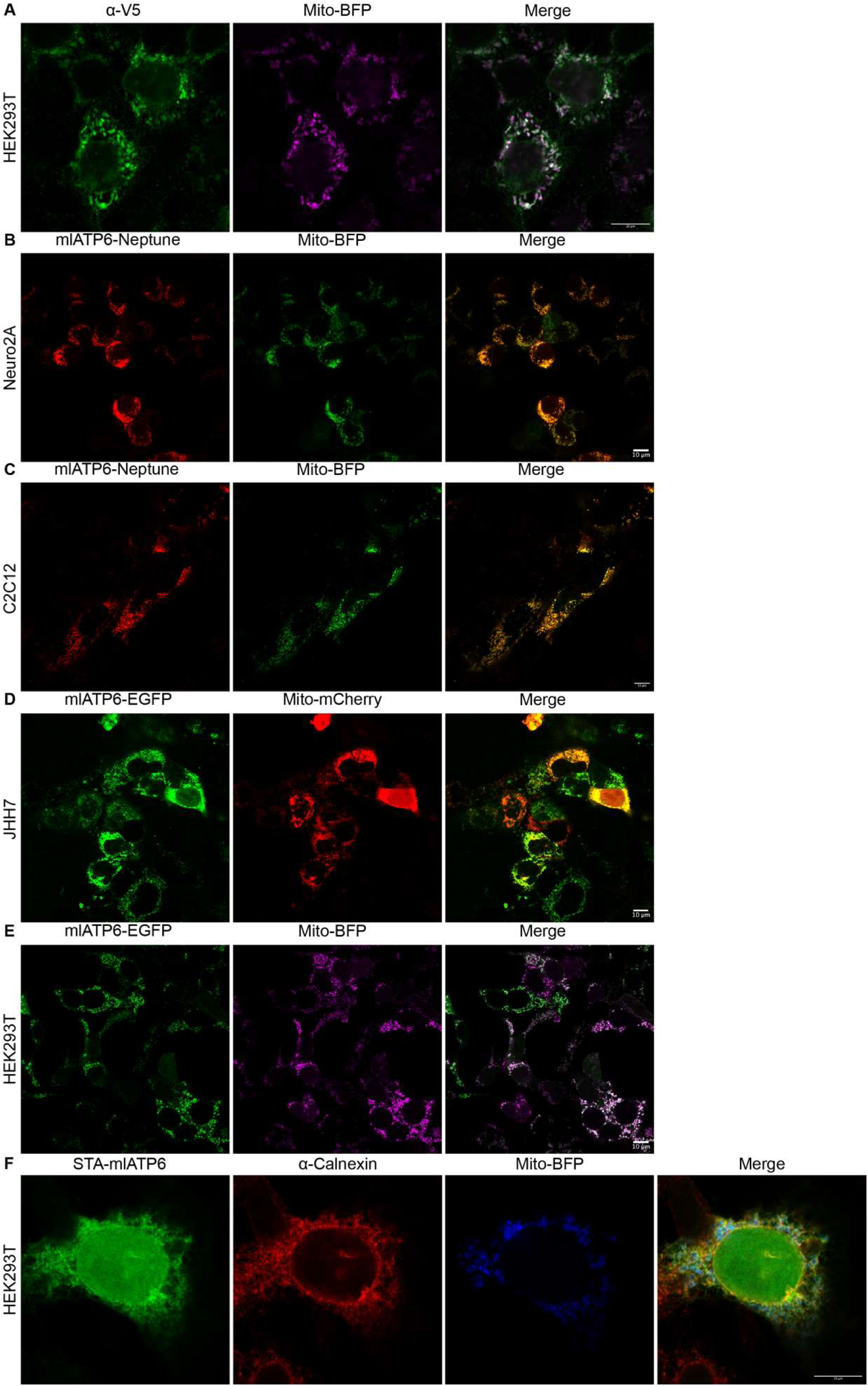
mlATP6 localizes to mitochondria across cell types. **(A)** HEK293T (human embryonic kidney) cells expressing mlATP6-V5, detected by anti-V5 immunofluorescence (green) and Mito-BFP (magenta). Merge shown at right. **(B)** Neuro2A (mouse neuroblastoma) cells expressing mlATP6-Neptune (red) and Mito-BFP (green). Merge shown at right. **(C)** C2C12 (mouse skeletal muscle) cells expressing mlATP6-Neptune (red) and Mito-BFP (green). Merge shown at right. **(D)** JHH7 (human hepatocellular carcinoma) cells expressing mlATP6-EGFP (green) and Mito-mCherry (red). Merge shown at right. **(E)** HEK293T cells expressing mlATP6-EGFP (green) and Mito-BFP (magenta). Merge shown at right. **(F)** HEK293T cells expressing STA-mlATP6-EGFP (green), stained for the ER marker calnexin (anti-calnexin, red), with Mito-BFP (blue). Merge shown at right. Scale bar, 10 µm.

**Figure 7—figure supplement 1:**
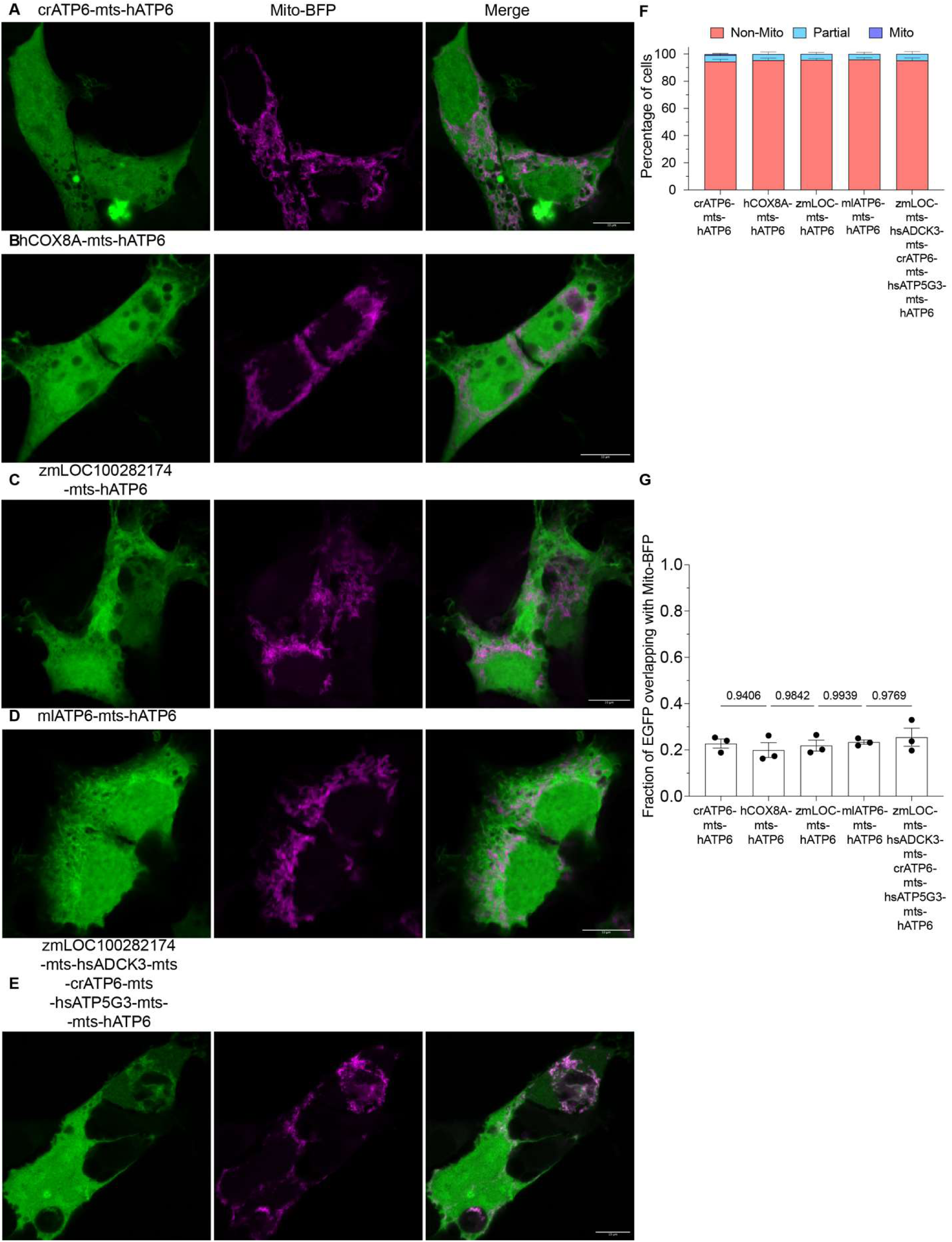
Strong MTSs alone are insufficient for hATP6 targeting. **(A–E)** Representative confocal images of HEK293T cells expressing the indicated MTS-hATP6 constructs (EGFP, green) and Mito-BFP (magenta). Scale bars, 10 µm. **(F)** Scoring of EGFP localization across cells as Non-Mito, Partial, or Mito for each construct. Data are mean ± SD from three independent experiments. Representative images of each scoring category (Non-Mito, Partial, Mito; left to right) are shown above. **(G)** Manders’ colocalization coefficient (M1; fraction of EGFP overlapping with Mito-BFP) quantifying mitochondrial localization for each construct. Data are mean ± SEM from three independent experiments; p-values from repeated-measures one-way ANOVA with Tukey’s multiple comparisons test.

**Figure 7—figure supplement 2:**
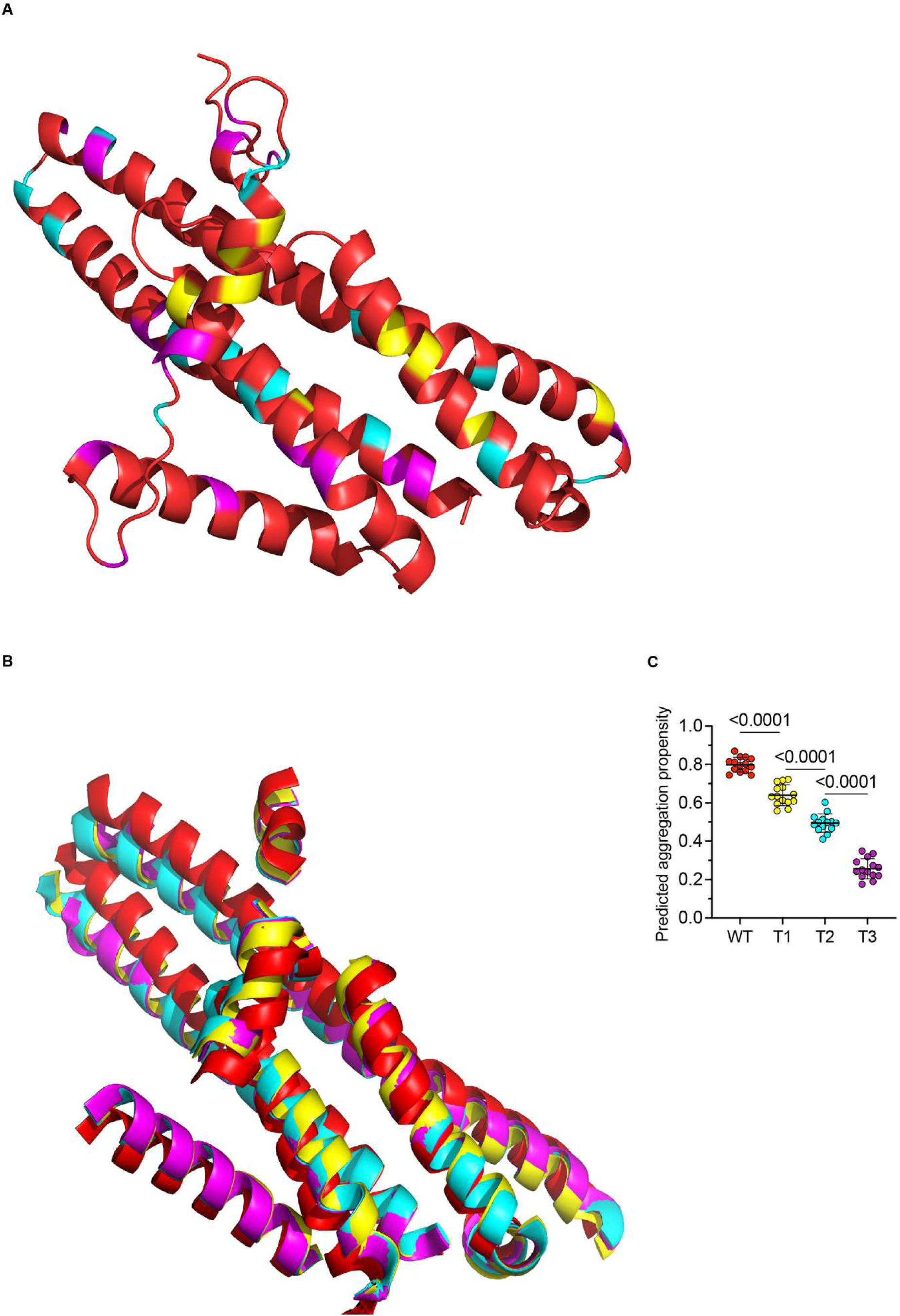
ATS substitutions preserve hATP6 structure. **(A)** AlphaFold2-predicted structure of hATP6-WT (red) with residues substituted in each ATS construct highlighted: T1 substitutions (yellow), T2 substitutions (cyan), T3 substitutions (magenta). **(B)** Structural alignment of independently predicted AlphaFold2 structures for hATP6-WT (red), T1 (yellow), T2 (cyan), and T3 (magenta), each colored as a whole structure. Predicted MTS and loops removed for clarity. **(C)** Predicted aggregation propensity scores for hATP6-WT, -T1, -T2, and -T3. Bars show mean ± SD. p-values from one-way ANOVA with Tukey’s post hoc test.

**Figure 7—figure supplement 3:**
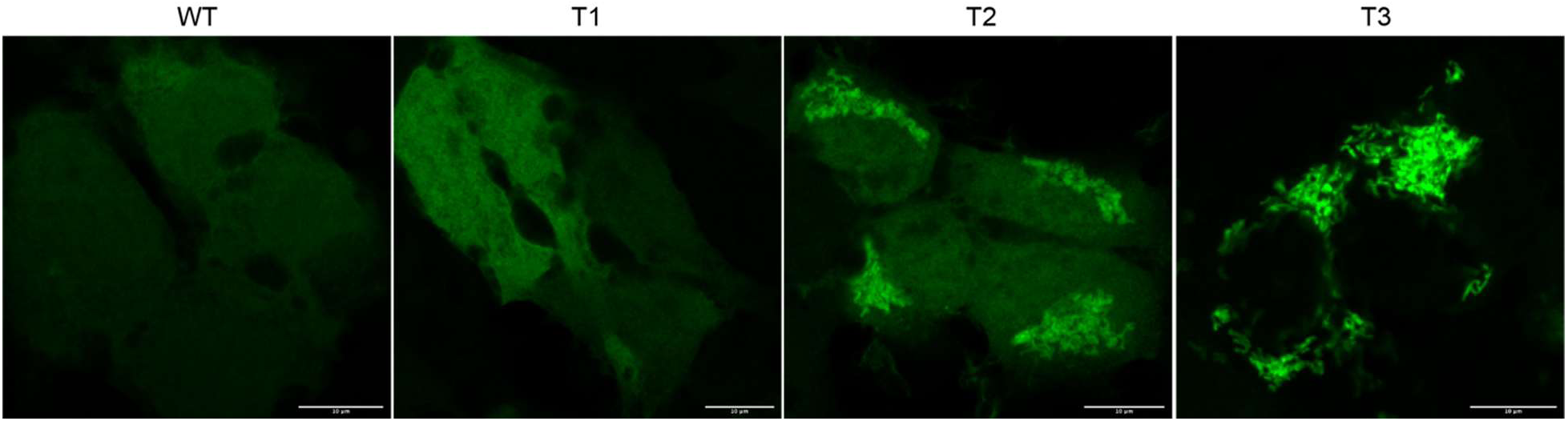
Targeting tracks with hydrophobicity, not expression. Representative confocal images of HEK293T cells stably expressing hATP6-WT, -T1, -T2, or -T3 (left to right; EGFP, green), acquired under identical confocal settings. Lower-expressing cells were selected for WT and T1; higher-expressing cells were selected for T2 and T3. Scale bars, 10 µm.

**Figure 8—figure supplement 1:**
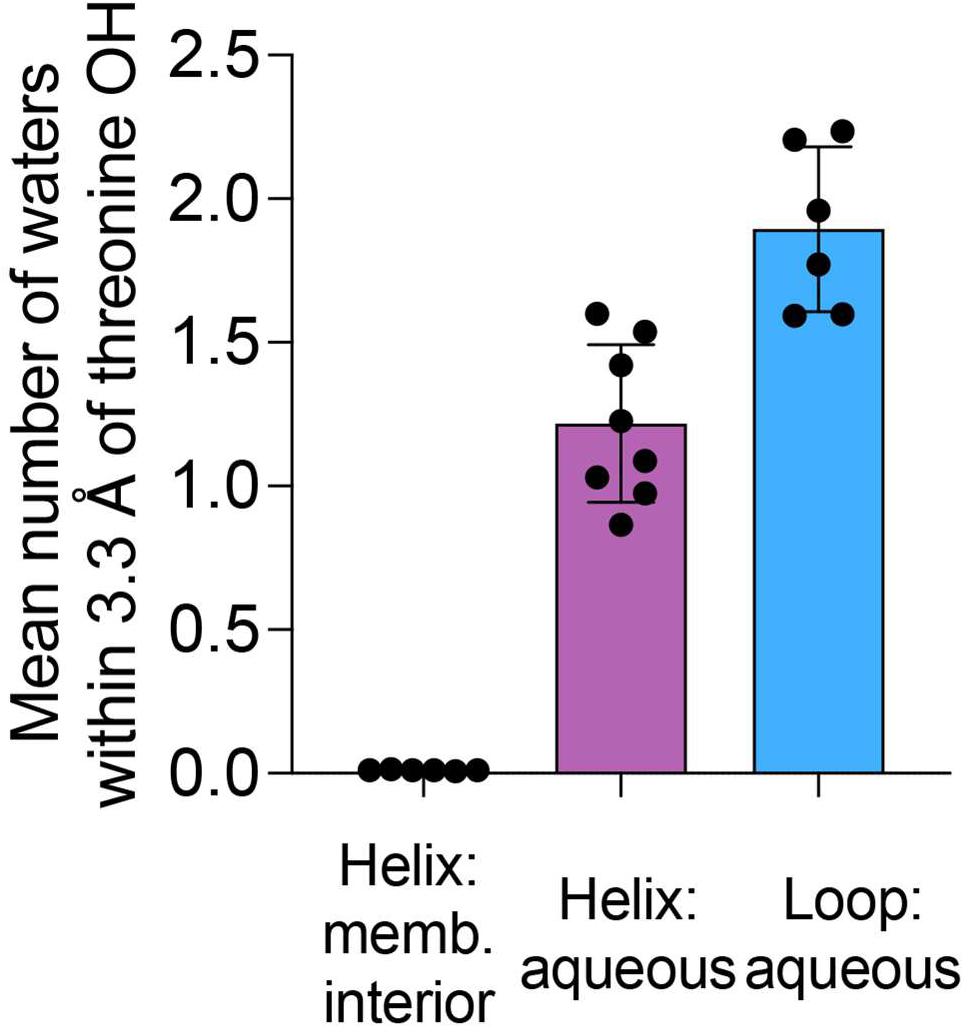
Threonine hydroxyl solvation by structural and solvent context. Mean number of water molecules within 3.3 Å of the threonine sidechain hydroxyl, for threonines classified as intrahelical/buried, intrahelical/aqueous, or loop/aqueous, averaged across all simulations. Error bars denote SD.

## METHODS

### Evolutionarily related protein pair identification

To infer evolutionary relationships between *E. coli* and *H. sapiens* proteomes, InParanoiDB v9 (accessed in April 2024) was queried using the species-pair search tool to find ortholog groups between *Escherichia coli* strain K12 (taxID: 83333) and *Homo sapiens* (taxID: 9606) (Persson and Sonnhammer, 2022). Domain orthologs from each orthology group were extracted as protein pairs linking *E. coli* proteins to related human proteins. If a single *E. coli* protein was associated with multiple distinct human proteins, each association was treated as an evolutionarily related protein pair. However, redundant associations involving alternative isoforms of the same human protein were retained as a single representative pair based on the highest inparalog score (an InParanoid sequence-similarity confidence score, typically corresponding to the canonical UniProtKB isoform). The resulting pairwise associations were subsequently categorized into transmembrane or non-transmembrane/ soluble groups according to the presence or absence of UniProtKB transmembrane-region annotations (UniProt Consortium, 2023). All UniProtKB accessions for both transmembrane and soluble protein pairs are provided in supplementary tables A and B.

### Structure-informed sequence alignment

AlphaFold2-predicted structures for evolutionarily related protein pairs were obtained as the default model provided for each UniProtKB accession (Jumper et al., 2021). For proteins without available entries, structures were predicted using ColabFold v1.6.1 in monomer mode with default parameters and the top-ranked model was used for downstream analyses (Mirdita et al., 2022).

Paired protein structures were structurally aligned using the *align* command in PyMOL. Alignments were restricted to corresponding helical regions, and only regions with clear one-to-one visual correspondence in the structure were retained. Helical regions were defined based on secondary-structure assignment from AlphaFold models in PyMOL, and boundaries were determined based on continuous α-helical segments.

These corresponding helical regions were mapped between proteins and number of transmembrane helices was verified using TMHMM 2.0 (Krogh et al., 2001). Transmembrane helices were defined based on agreement with UniProtKB annotations and TMHMM 2.0 predictions where UniProtKB annotations were not available. The boundaries from these annotations were refined using AlphaFold2 structures to include the full extent of each continuous α-helical stretch. Structural context, such as helix packing and orientation, was used to confirm membrane-spanning geometry and only helices consistent with canonical transmembrane architecture (length ≥ 15 residues and parallel stacking architecture) were retained. Helices that shared no overlap with those classified as transmembrane were designated as soluble. This category comprises all α-helical regions that showed correspondence between evolutionarily related proteins, including extramembrane and potentially amphipathic helices, without additional filtering criteria. Both mitochondrial and non-mitochondrial soluble protein groups were processed similarly.

Paired helix sequences were aligned in R using the *pairwiseAlignment* function with PAM250 matrix; a gap opening penalty of 20 and a gap extension penalty of 20. The same alignment parameters were applied uniformly across all protein groups, including transmembrane, soluble, mitochondrial, and non-mitochondrial proteins.

### Hydrophobicity calculation

The GRAVY hydrophobicity score was calculated using the *hydrophobicity* function from the *Peptides* package in R, based on the Kyte-Doolittle scale (Kyte and Doolittle, 1982). For evolutionarily related protein pair analyses, unaligned helix sequences were used for hydrophobicity comparison between paired helices. For whole protein hydrophobicity analyses, MTS, epitope tags and linker sequences were removed prior to hydrophobicity comparison across protein constructs.

### Amino acid composition analysis

Amino acid composition was calculated for each helix as the fraction of residues corresponding to predefined categories based on biochemical properties or individual amino acids of interest. In particular, the fraction of aliphatic residues (A, L, I, V, and F), threonine, and serine was quantified for each transmembrane or soluble helix. For pairwise analyses, matched helices from aligned protein pairs were compared directly on a one-to-one basis as these represent distinct structural elements. These paired comparisons were used for downstream statistical testing.

### Substitution matrix analysis

Amino acid substitution frequencies were calculated from aligned residues within transmembrane helices from evolutionarily related protein pairs. For each aligned residue pair from these sequence pairs, the *E. coli* residue was assigned to the matrix row, and the human residue was assigned to the matrix column. Thus, substitution matrix entries represent observed *E. coli* → human residue pairings across aligned helix positions. Gapped alignment positions were excluded, and identical residue pairs representing residue conservations were retained as diagonal entries.

For each pair of helices being compared, counts were tabulated for all residue-residue pairings observed across aligned positions. Substitution matrices were generated separately for each protein pair and combined by element-wise summation for specified protein groups. Given the small number of mtDNA-encoded IMM proteins, nuclear- and mtDNA-encoded IMM proteins were pooled into a single mitochondrial group, reflecting their shared membrane environment and aggregation constraints. Equivalent procedures were applied to control groups of soluble helices where indicated. For relative substitution matrix generation, individual substitution count matrices were normalized by row sums, followed by element-wise division of the normalized mitochondrial matrix by the normalized non-mitochondrial control matrix. No pseudocounts were added; entries with zero denominator values were set to NA before visualization. Substitution matrices were visualized using the *ComplexHeatmap* package in R.

Statistical analyses for enrichment were carried out for each substitution using Fisher’s exact test (two-sided). For each E. coli residue *i* and human residue *j*, the contingency table compared the number of *i*→*j* pairings in the test set with all other pairings from the same E. coli residue in the test set, against the corresponding counts in the control set. Unadjusted p-values are reported for individual substitutions and interpreted for enrichment patterns.

### Positional analysis across transmembrane helices

To analyze residue distributions across transmembrane helices, sequences annotated in UniProtKB were aligned using a common positional framework centered on the helix midpoint.

In topology-agnostic analyses, helices were oriented without regard to membrane sidedness. Positional profiles were then symmetrized by averaging frequencies at equivalent positions on either side of the helix midpoint; for example, positions +3 and −3 were assigned the same averaged value.

For topology-informed analyses, helix orientation according to cytosolic→matrix/ luminal direction was assigned using the OPM database and verified against PDB structural models (Lomize et al., 2006). Where structural orientation was not available in the OPM database, membrane topology was assigned based on published literature. Only proteins with experimentally supported or explicitly reported orientation were retained.

For both analyses, residue frequencies were computed at each relative position across the aligned helices. The central 11 positions were defined as the core, while the five positions on either side were designated as peripheral regions. For helices longer than 21 residues, residues outside the central 21-position window were excluded from the analysis. For helices shorter than 21 residues, only observed residues were included at their corresponding relative positions, and missing terminal positions were not imputed or counted as zeros. Enrichment in core relative to peripheral regions was assessed using a two-sided Fisher’s exact test in GraphPad QuickCalcs (GraphPad Software, www.graphpad.com, accessed January 2026). For positional enrichment plots, residue fraction line plots were smoothed using a rolling average with a window size of 5 and symmetrized about the central position and visualized using GraphPad Prism v10.5. Smoothing and symmetrization were applied only to residue-fraction line plots for visualization and were not used for statistical testing.

### Proteome-wide compartment comparisons

For the human proteome, transmembrane proteins were grouped based on UniProtKB-annotated subcellular localization and membrane classification. Protein sets were generated using predefined UniProtKB query filters and were not additionally filtered by protein length, transmembrane helix number or AlphaFold confidence. The exact queries and the UniProtKB-accessions for proteins from each compartment are provided in supplementary table C. For ER and Golgi comparisons, proteins were selected when annotated to the specified compartment and when structural information was available. For the nuclear group, 41 proteins met the criteria of being transmembrane and nuclear annotated in UniProtKB and were included in the analysis. All transmembrane proteins annotated as outer mitochondrial membrane or inner mitochondrial membrane were included in the mitochondrial analyses. Proteins annotated to unrelated organelle categories were excluded from compartment-specific sets.

Although experimental structures were used as a selection criterion, AlphaFold-predicted structures were used for downstream compositional analyses to ensure complete protein coverage. Experimental structure availability was used only to select proteins with prior structural support.

To analyze the broader human transmembrane proteome, reviewed human UniProtKB / Swiss-Prot entries annotated with transmembrane helices were retrieved from UniProtKB. Proteins were retained if they contained a UniProtKB-defined transmembrane helix annotation and a curated subcellular-location annotation. The resulting set of reviewed human transmembrane proteins was used for subsequent compositional analyses.

For each protein, residue composition was calculated by concatenating all transmembrane helices or all soluble helices, yielding a single per-protein estimate of residue frequency. For analyses comparing residue distributions within helices (e.g., threonine versus serine), calculations were performed at the individual helix level.

For heatmap generation, relative enrichment was calculated as the fold change in mean residue fractions relative to a non-mitochondrial reference group. To prevent compartments with more proteins from skewing the reference, the non-mitochondrial reference was defined as the unweighted mean across endoplasmic reticulum, Golgi, cell membrane, and nuclear residue fractions.

### Metazoan and Plantae TMH dataset construction

Protein sequences and annotations were obtained from UniProtKB (accessed January 2026). For metazoan TMH analyses, UniProtKB was queried across Metazoa, and only species with proteins annotated as mitochondrial genome–encoded in UniProtKB were retained for downstream cross-compartment comparisons.

For Plantae proteins, UniProtKB was queried across Viridiplantae, and only species with both chloroplast genome-encoded and mitochondrial genome-encoded proteins annotated in UniProtKB were retained.

For both datasets, redundant homologous proteins were removed using CD-HIT with a 90% sequence identity threshold (Fu et al., 2012). Proteins from the selected species were pooled only after this filtering step, and all TMHs from each protein were concatenated to generate a single per-protein sequence for downstream compositional analyses.

### Soluble helix controls from TM proteins

As an internal control, soluble (extramembrane) helices were extracted from proteins that also contained transmembrane helices and analyzed separately from membrane-spanning segments. All helical regions were extracted from AlphaFold-predicted models as visualized in PyMOL and then transmembrane helices were excluded. Transmembrane helices were initially identified based on UniProtKB annotations and TMHMM 2.0 predictions, and boundaries were refined using AlphaFold structural models to include the full extent of each continuous helix. Helices with any overlap with these refined transmembrane regions were excluded from the soluble control set. Soluble helices were therefore defined as α-helical regions that did not overlap any transmembrane helix. All soluble helices from each protein were concatenated to generate a single per-protein sequence, ensuring that proteins containing multiple helices did not contribute disproportionately to subsequent analyses. This definition includes all non-transmembrane α-helical regions (e.g. signal peptides, targeting sequences, or amphipathic helices) – without additional filtering. As mtDNA-encoded TM proteins possess only a few soluble helices, both concatenated and non-concatenated comparisons are provided for helix-level analyses from this set.

Proteins lacking any predicted transmembrane segments were treated as soluble proteins and all helical regions were concatenated for compositional analyses. Hydrophobicity and residue composition were calculated using the same procedures applied to TMHs. These analyses were performed within the same protein set used for transmembrane helix comparisons. No further filtering was applied.

### Analysis of mtDNA-to-nuDNA gene transfer events

Protein sequences of mtDNA-encoded proteins from *Andalucia godoyi* were obtained from UniProtKB (accessed August 2024) as per the list reported by Gray et al. (Gray et al., 2020). The SecY protein sequence was obtained from the related jakobid *Reclinomonas americana*. All *A. godoyi* proteins were queried using BLASTp against proteins from eukaryotic phyla in which the corresponding gene has been reported as absent from the mitochondrial genome by Butenko et al. (Butenko et al., 2024). No fixed sequence identity or coverage thresholds were applied at this stage. Instead, resulting candidate hits were grouped as per taxonomic classification and representative protein sequences from each taxonomic lineage were examined to assess nuclear encoding.

Successful nuclear transfer was inferred based on the presence of an N-terminal MTS using TargetP 2.0 prediction based on a probability cutoff of 0.4 (Almagro Armenteros et al., 2019). Proteins that showed dual targeting to both chloroplast and mitochondria were excluded from further analysis. Nuclear expression was also confirmed by assessing the expression and genomic localization of candidate protein in a nuclear RNA or DNA sequencing dataset, wherever such datasets were available. For each successful instance of nuclear transfer, a closely related lineage retaining the mtDNA-encoded version of the protein was identified. Mitochondrial genome encoding in this lineage was assigned based on UniProtKB annotation or corresponding mitochondrial genome records. Absence of a TargetP 2.0-predicted MTS and sequence similarity to *A. godoyi* reference and nuDNA-encoded protein, as assessed using CLUSTAL W, were used as supporting criteria to confirm mitochondrial encoding.

For each lineage pair, the species with the highest pairwise cytochrome b identity (assessed by BLASTp) was selected as the representative for downstream analyses. Being a core subunit of the mitochondrial respiratory chain and universally encoded by mtDNA, cytochrome b is an operational proxy for evolutionary relatedness among mitochondrial genomes. Accession identifiers for all analyzed proteins are provided in supplementary table D.

Structural alignment was then performed between AlphaFold structures using PyMOL for mtDNA- and nuDNA-encoded proteins from each pair. Subsequently, compositional analyses were carried out on aligned transmembrane and soluble helix sequences from each protein pair as described previously.

### Genome-wide TMH composition control

To control for potential differences in genome-wide TMH composition across our phylogenetically diverse organismal pairs, we calculated a per-organism threonine asymmetry score. For each organism, we computed the mean threonine content of all UniProtKB-annotated nuDNA-encoded TM proteins divided by the mean threonine content of all mtDNA-encoded TM proteins. Per-protein threonine fractions were calculated from concatenated TMH sequences as described above. The resulting ratios were compared between mtDNA-encoding (n=12) and nuDNA-encoding (n=12) organisms — paired by our organismal pair set — using Wilcoxon signed-rank test.

### mtDNA-to-nuDNA residue asymmetry analysis

Soluble and transmembrane proteins that underwent gene transfers were structurally aligned to extract sequences for analogous soluble and transmembrane helices between mtDNA-encoded and nuDNA-encoded proteins. Paired sequences were aligned using the *pairwiseAlignment* function in R with BLOSUM62 substitution matrix with gap opening penalty of 20 and gap extension penalty of 5. The default BLOSUM62 was used for these alignments because mtDNA-encoded and nuDNA-encoded homologs from closely related lineages were substantially more similar than the more divergent E. coli–human protein pairs analyzed above. These parameters were kept consistent for aligning paired helix sequences from transmembrane and soluble proteins. Resulting counts of substitutions were collected from alignment between mtDNA-encoded and nuDNA-encoded helix sequences and compiled into raw substitution matrices.

To assess whether substitutions originating from aliphatic residues (ALIVF) to a particular target amino acid were over-represented, we performed a directional enrichment analysis. For each residue X, the number of substitutions from aliphatic residues to X (ALIVF→X) and the number of substitutions from X to aliphatic residues (X→ALIVF) were calculated. Corresponding directional counts for background substitutions were obtained by considering substitutions involving all residues minus those defined as aliphatic residues. For each target amino acid, a 2×2 contingency table was constructed using substitutions from aliphatic versus non-aliphatic residues to X and from X to aliphatic versus non-aliphatic residues.

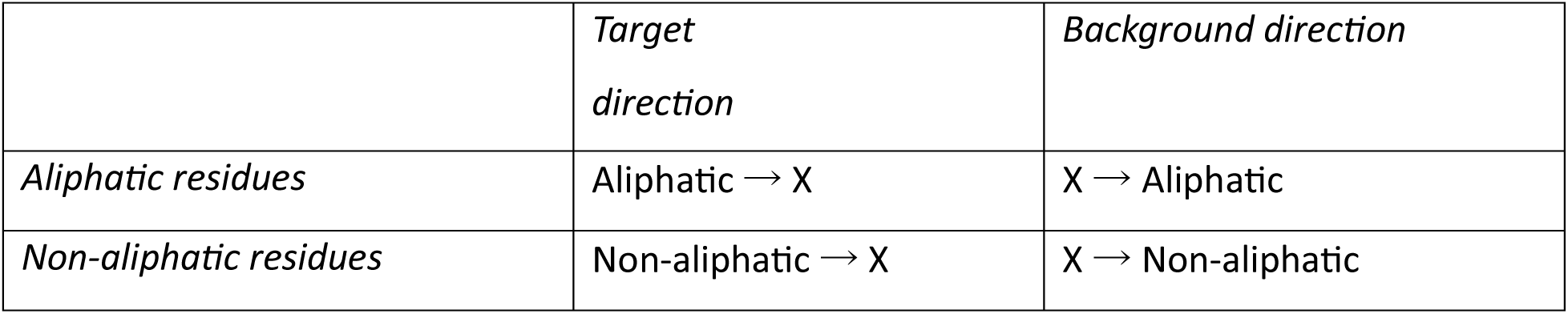

Statistical significance of enrichment was assessed using Fisher’s exact test with a one-sided alternative hypothesis (greater) in R. p-values were adjusted for multiple testing across all amino acids using the Benjamini-Hochberg false discovery rate (FDR) correction.

### Predicted aggregation propensity analysis

Predicted aggregation propensity was estimated using Aggrescan3D 2.0 (accessed in December 2025) (Pujols et al., 2022). AlphaFold-predicted monomeric structures were used as input and structures were energy-minimized prior to scoring using Aggrescan3D’s built-in minimizer. Stability calculations were performed in native structural mode (without mutation scanning) with an aggregation sphere radius of 10 Å.

For gene transfer comparisons between naturally occurring proteins, dynamic mode was turned off and global average aggregation propensity scores from static input structures were compared for protein pairs. MTS regions were retained in the primary analysis because they represent native sequence features of nuDNA-encoded proteins. Analyses repeated after removal of predicted MTS regions produced similar results and are not shown.

For engineered hATP6 constructs, MTS sequences, linkers, and tags were removed before AlphaFold modeling and Aggrescan3D analysis, such that only the ATP6 coding sequence was analyzed. Both static and dynamic Aggrescan3D analyses were performed as dynamic-mode scores account for potential conformational flexibility introduced by ATS mutations. Both modes yielded similar results.

### Phylogenetic analysis of ATP6 and ATP9

ATP6 and ATP9 sequences were collected from representative taxa containing mitochondrial-encoded or nuclear-encoded variants. For nuDNA-encoded sequences, MTS regions were removed before further analyses based on TargetP 2.0 predicted cleavage positions. Multiple sequence alignments were generated using MAFFT (v7.511) with the L-INS-I algorithm and default parameters (Katoh and Standley, 2013). Phylogenetic trees were generated using Neighbor-Joining method, with the JTT substitution model and no rate heterogeneity and default parameters. Branch support estimation was performed using 1000 bootstrap replicates. Phylogenetic trees were visualized using Phylo.io (Robinson et al., 2016).

### Identification of mlATP6 sequence

*Mnemiopsis leidyi* genome was downloaded from GenBank (Accession GCA_000226015.1). The *M. leidyi* genome was queried using tBLASTn with ATP6 from *Aurelia limbata* (UniProtKB: A0A6G6CGA7) as the query. The resulting mlATP6 hit was then used to query the EST dataset to identify supporting transcript evidence. Multiple independent EST matches (16 total) were used to provide transcript support and refine the mlATP6 coding sequence. Supporting EST sequences are provided in supplementary table E. The resulting mlATP6 sequence was further validated based on high (82.87%) identity with ATP6 from another ctenophore, *Pleurobrachia bachei* (Kohn et al., 2012).

### Design and expression of mlATP6 constructs

The nuDNA-encoded mlATP6 coding sequence of comb jelly *M. leidyi* obtained from earlier analyses was aligned with the mtDNA-encoded ATP6 sequence of the brown moon jelly *Aurelia limbata* (al) using CLUSTAL W. Substitutions of threonine to aliphatic residues (STA) were made between mlATP6 and alATP6 to generate the mlATP6-STA construct, which retained the native mlATP6 MTS. The human MT-ATP6 sequence was designated as the hATP6-WT construct and appended with an mlATP6 MTS on the N-terminus following sequence alignment using CLUSTAL W.

All three sequences — mlATP6-WT, mlATP6-STA, and hATP6-WT, along with the substitution list, are provided in supplementary table F. All constructs were recoded for human nuclear expression and codon optimized using IDT codon-optimization tool (www.idtdna.com). Constructs were cloned into pLentiBlast vector under the CMV promoter and with a C-terminal EGFP tag joined by an -SSG linker (https://www.addgene.org/125133/). For mlATP6-WT constructs, constructs encoding other C-terminal tags (FLAG and V5) were also generated using the same procedure. The mlATP6-Neptune construct was generated by cloning the codon-optimized coding sequence into pmNeptune2-N1 vector upstream of mNeptune2.

For expression, HEK293T cells were transfected with plasmids using Lipofectamine 2000 and labeled with MitoTracker Red 24 hours post-transfection. Subcellular localization was assessed using fluorescence microscopy as described below.

mlATP6-WT-EGFP construct cloned into pLentiBlast was also used to generate lentiviral particles for transduction experiments. Lentiviral particles were produced in HEK293T cells using the EPFL 293T transient-transfection protocol.

### Design and expression of hATP6-WT constructs with various MTSs

MTS sequences were obtained from published studies reporting targeting of hATP6 to the mitochondria and are provided in supplementary table G. These sequences were appended to the N-terminus of human MT-ATP6 and EGFP tagged on the C-terminus via an SSG linker in a pLentiBlast backbone under the CMV promoter (MTS–hATP6–SSG–EGFP). All constructs were codon-optimized for nuclear expression and verified using Sanger sequencing.

For expression, HEK293T cells at ∼70% confluency were transfected with the above plasmid constructs using Lipofectamine 2000 (Thermo Fisher; Cat# 11668027) according to the manufacturer’s protocol. Cells were imaged 24 hours post-transfection for subcellular localization (see fluorescence microscopy methods below).

### Design and expression of hATP6-ATS constructs

The human MT-ATP6 sequence was compared with 16 other mitochondrially-encoded ATP6s-15 from metazoan species and 1 from *Saccharomyces cerevisiae* (ATP6 sequence accession identifiers and CLUSTAL W multiple sequence alignment provided in supplementary table H).

A series of ATS constructs (T1, T2, T3) was made from the hATP6-WT sequence by substituting aliphatic residues to threonine based on their conservation in the alignment.

Substitutions were introduced in a stepwise manner: T1 included substitutions at positions where threonine is commonly observed in the alignment, and the aliphatic residues are not highly conserved; whereas T2 and T3 incorporated additional ATS swaps where threonines occur less frequently in the alignment. Consequently, the number of ATS swaps increases progressively from T1 to T3, reflecting an increasing extent of threonine incorporation. The aligned construct sequences are provided in supplementary table I.

For example, the alanine at position 46 aligned to a threonine in multiple ATP6 sequences and was therefore substituted in T1. In contrast, valine at position 56 aligned to threonine in a single ATP6 and was substituted only in the T2 and T3 constructs.

Following ATS swaps, the mlATP6 MTS sequence was appended to the N-terminus of all hATP6-ATS constructs and codon optimization was carried out. All constructs were cloned into the pLentiBlast vector backbone with a C-terminal EGFP tag fused with an -SSG linker and lentiviral particles were generated.

For expression, HEK293T cells were transduced with lentiviral particles encoding hATP6-ATS-EGFP constructs. Culture medium was first replaced with fresh medium containing 1 µg/mL polybrene (Sigma-Aldrich, Cat# TR-1003), and cells were incubated for 20 min at 37 °C. Lentiviral supernatant was then added at a 1:1 ratio with culture medium. After 24 h, cells were selected with 10 µg/mL blasticidin (Gibco, Cat# A1113903) for 3 days, until cells in untransduced control wells were no longer viable. Following selection, cells were maintained in culture medium without blasticidin. For assessing colocalization, cells were grown on coverslips, labeled with MitoTracker Red and DAPI and imaged as described below.

### Cloning

Constructs were cloned using *E. coli* NEB Stable competent cells (New England Biolabs) followed by transformation according to manufacturer instructions and grown at 30 °C for lentiviral constructs and 37 °C for non-lentiviral constructs overnight. All constructs were generated by restriction enzyme-based cloning using PCR-amplified inserts and verified by Sanger sequencing across the full coding region.

### Cell culture and expression conditions

The following mammalian cell lines were obtained: HEK293T (ATCC, CRL-11268), Neuro2A (ATCC, CCL-131), C2C12 (ATCC, CRL-1772) and JHH7 (JCRB Cell Bank, JCRB1031). Cells were used between passages 5 and 20. All mammalian cell lines used for our experiments were maintained under identical conditions at 37 ℃ at 5% CO_2_ in DMEM (Corning, Cat# 10-013-CV) supplemented with 10% Fetal Bovine Serum (GenClone, Cat# 25-550H) and 1% penicillin-streptomycin (1000 U/mL; Gibco, Cat# 15140122). Cultures were routinely screened for mycoplasma by DAPI staining. For transfection experiments, cells were seeded in 12-well plates to achieve ∼70% confluency and transferred to coverslips for imaging after 24 h.

### Fluorescence Microscopy

For EGFP-tagged constructs, EGFP fluorescence was imaged directly. For -V5 and -FLAG immunostaining, cells were plated on glass coverslips with poly-D-lysine and laminin coating (Neuvitro, Cat# GG-18-Laminin) and tagged constructs were expressed as specified in their respective methods sections. After 24 h, cells were treated with 20 nM MitoTracker Red CMXRos (Invitrogen, Cat# M46752) for 30 min at 37 ℃ and fixed with 4% paraformaldehyde in PBS for 15 min. After 3 PBS washes, fixed samples were permeabilized with 0.1% Triton X-100 in PBS for 15 min and washed 3 times with PBS. Blocking with a solution of 5% Normal Goat Serum, 1% Bovine Serum Albumin and 0.05% Tween-20 in PBS was performed for 1 h at room temperature. Immunofluorescence was performed using Anti-V5 (Bio-Rad, Cat# MCA1360) or Anti-FLAG (Sigma-Aldrich, Cat# F3165-1MG) primary antibodies diluted 1:1000 in fresh blocking solution at 4 ℃ overnight. Coverslips were washed thrice with PBS and incubated with Alexa Fluor 488 goat anti-mouse secondary antibody (Invitrogen, Cat# A-11001) prepared in blocking solution for 1 h at room temperature. DAPI (Sigma-Aldrich, Cat# D9542) staining was performed for 15 min at room temperature with 1 μg/mL solution in PBS and coverslips were washed four times with PBS. Coverslips were mounted with Vectashield Vibrance antifade mounting medium (Vector Laboratories, H-1700) and imaging was performed on a Zeiss LSM 880 confocal microscope with a 63x oil-immersion objective (NA 1.4). Fluorophores were excited using 405 nm (DAPI), 488 nm (EGFP/ Alexa Fluor 488), and 561 nm (MitoTracker Red) lasers and channels were acquired sequentially to avoid bleed-through. Images were acquired as single optical sections with pinhole size set to 1 Airy unit. Laser power was kept constant across samples within each colocalization experiment, while detector gain was adjusted as needed to optimize signal detection and avoid saturation. For experiments involving quantitative intensity comparisons, all acquisition settings, including laser power and detector gain, were kept constant across samples. Raw images were processed using FIJI.

### Localization scoring and colocalization quantification

Scoring for localization was performed using manual phenotype scoring with blinding to experimental construct. Cells were scored into three categories: A, non-mitochondrial (includes diffuse/cytosolic localization); B, partially mitochondrial with cytosolic signal; and C, predominantly mitochondrial with minimal cytosolic signal. All experiments were repeated for three biological replicates and at least 50 cells per replicate per condition were imaged for scoring. Scoring criteria were defined prior to analysis and kept consistent throughout all conditions across all experiments. To avoid potential bleed-through from the DAPI or MitoTracker Red channels affecting manual localization scoring, cells from the same polyclonal population were seeded on parallel coverslips and imaged without these dyes for scoring; dye-stained coverslips were imaged separately for representative images and colocalization analysis.

For colocalization analysis, at least 15 cells were imaged per replicate per condition. To ensure representation across localization phenotypes, cells were sampled from each category (A,B,C) for high-resolution imaging. Regions of interest were defined by selecting entire cells. Colocalization between construct signal and MitoTracker was quantified using the JACoP plugin to calculate Pearson’s r and Manders’ overlap coefficients (M1 and M2). All images were identically processed; no smoothing or background subtraction was applied to GFP channel, in which all our constructs were imaged. All thresholding parameters (manual thresholding) were kept constant across all constructs throughout all experiments.

### MD system construction

We performed a new analysis of MD simulation trajectories published in Borowsky and Grabe (Borowsky and Grabe, 2025). The protocol is outlined below for reference.

MD simulations were performed on ATP/ADP Carrier Protein 1 (ANT1, PDB: 2C3E) and Uncoupling Protein 1 (UCP1, PDB: 8HBV). Each protein (and its three associated cardiolipins) was embedded in a POPC bilayer with 7 mol% arachidonic acid and solvated with TIP3P water using the CHARMM-GUI membrane builder. Both proteins were parameterized using the CHARMM36m force field, and lipids were parameterized using CHARMM36. All-atom MD simulations were carried out using GROMACS 2024. Systems were simulated at 310.15 K over six independent replicates, for an aggregate simulation time of 75 μs and an average molecular time of 6 μs.

### Residue selection and helical categorization

Residue-level secondary structure assignment was performed using the MDAnalysis DSSP tool. Threonine residues in each frame were then categorized as helical if the four preceding residues were assigned as helical by DSSP to guarantee the availability of a backbone carbonyl at the *i−4* position for hydrogen-bonding with threonine. Only residues categorized as helical for ≥ 75% of the total simulation frames were classified as helical for the purposes of further analysis. Threonine residues were classified as being on loops if they were assigned as being in the loop state by DSSP for ≥ 75% of the total simulation frames. No residues were classified as being on both loops and helices because of the way the DSSP assignments are defined from hydrogen bond contact maps and the geometric constraints associated with the preceding four residues having helical H-bonding patterns.

### Solvation environment classification

The solvent environment of each helical residue was quantified by calculating the average number of hydrogen bonds formed between the sidechain hydroxyl group and water molecules. Using MDAnalysis HydrogenBondAnalysis, we established discrete solvation categories:

Intramembrane (mem): < 0.05 hydrogen bonds per frame

Interfacial: 0.05 ≤ value < 0.67 hydrogen bonds per frame

Aqueous (aq): ≥ 0.67 hydrogen bonds per frame

These thresholds were used to isolate the energetic impact of the hydrophobic membrane core versus the aqueous solvent.

To further examine the solvation patterns, we calculated the average number of water molecules within 3.3 Å of each threonine OH (by oxygen-oxygen distance) for residues in different solvent and secondary structure environments. The results, depicted in Figure 8—figure supplement 1, show that the number of nearby waters is consistent with the hydrogen-bond-based solvation categories.

### Free energy calculations from rotameric populations

Sidechain **χ**_1_ dihedral angles (centered at -60°, +60°, and +180°) were monitored for each residue. The ±60° rotamers were treated as a single configurational state representing the state capable of forming backbone hydrogen bonds, while the +180° rotamer represented the non-bonding state. The free energy of the +180° rotamer (***Δ****G_rot_*_|*i*_) for each residue *i* was calculated as:

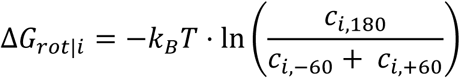

where *c* is the frame count for each state.

The effect of membrane insertion on the free energy of flipping from the ±60° rotamers to the 180° rotamer is ΔΔ*G* = Δ*G*_rot|mem_ − Δ*G*_rot|aq_ where *mem* and *aq* denote intramembrane and aqueous states respectively. This ΔΔ*G* value was calculated from simulations by averaging Δ*G*_rot|i_across aqueous or intramembrane residues to obtain Δ*G*_rot|aq_ and Δ*G*_rot|mem_respectively.

### Thermodynamic cycle construction for free energy estimation

The ΔΔ*G* introduced above describes a thermodynamic cycle with four states for each pair of solvation environment (aqueous, intramembrane) and rotamer category (±60°, 180°). ΔΔ*G* can therefore be defined using the membrane insertion legs of the cycle (with free energy ***Δ***G_ins_) rather than the rotamer flipping ones: ΔΔ*G* = Δ*G*_ins|180_ − Δ*G*_ins|±60_. The mathematical details of this are described in supplementary section 1.

MacCallum et al. (MacCallum et al., 2008) estimate the free energy of bringing amino acid side chains from water into the core of a phospholipid membrane using MD simulations. They use ethanol as a proxy for threonine (and methanol for serine), which is a good representation of the 180-degree rotamer which cannot hydrogen bond to the backbone, but not of the other two rotamers. Their insertion free energy ΔG_ins | Ethanol_ (or ΔG_ins | Methanol_) is therefore an estimate of ΔG_ins | 180_.

Hessa et. al. (Hessa et al., 2005) experimentally estimate the free energy of inserting threonine (and serine) on alpha helices from water into the membrane core (see their supplementary data S2 for the values). This is the overall free energy difference between the set of all aqueous rotamers and the set of all membrane-facing rotamers and is termed ΔG_ins_. In the limit where the ±60° rotamer is much more common than the 180° one, as is the case for threonine in both our MD simulations and crystallographic studies (Chamberlain and Bowie, 2002) the insertion energy is mainly determined by the former, so Δ*G*_ins|±60_ ≈ Δ*G*_ins_

### Membrane-water transfer energies and helical propensities of residues

Residue-specific membrane-insertion free energies for residues were taken from the experimentally derived scale of (Hessa et al., 2005). Helix–coil free-energy values, reflecting intrinsic α-helical propensities, were taken from Pace and Scholtz (Pace and Scholtz, 1998).

### Statistical analysis

All analyses were performed using GraphPad Prism 10 or R v4.5.2. Wilcoxon’s signed rank tests were used for pairwise comparisons, one-way ANOVA followed by Tukey’s post-hoc test was used for multiple comparisons. Unless otherwise specified, statistical analyses were performed using biologically independent units appropriate to each dataset. For helix-based analyses, individual helices were treated as observations unless aggregated at the protein level. For protein-level analyses, values were calculated per protein.

For cell-based experiments, statistical tests were performed on biological replicate means rather than individual cells and data are presented as Mean ± S.D from three independent biological replicates. For all other datasets, the ‘n’ and error bar definitions are indicated in the respective figure legends. Statistical significance was defined as p < 0.05. Custom R scripts used for all computational analyses will be deposited in a public repository upon publication. MD analysis scripts are available on GitHub at https://github.com/JonathanHB/threonine_rotamer_analysis.git.

